# Insights into the role of lipoteichoic acids and MprF function in *Bacillus subtilis*

**DOI:** 10.1101/2021.12.12.472321

**Authors:** Aurélie Guyet, Amirah Alofi, Richard A. Daniel

## Abstract

Gram-positive bacterial cells are protected from the environment by a cell envelope which comprises of layers of peptidoglycan that maintain the cell shape and teichoic acids polymers whose biological function remains unclear. In *Bacillus subtilis*, loss of all Class A Penicillin-Binding Proteins (aPBPs) which function in peptidoglycan synthesis is conditionally lethal. Here we show that this lethality is associated with an alteration of the lipoteichoic acids (LTA) and the accumulation of the major autolysin LytE in the cell wall. Our analysis provides further evidence that the length and abundance of LTA acts to regulate the cellular level and activity of autolytic enzymes, specifically LytE. Importantly, we identify a novel function for the aminoacyl-phosphatidylglycerol synthase MprF in the modulation of LTA biosynthesis in *B. subtilis* and *Staphylococcus aureus*. This finding has implications for our understanding of antimicrobial resistance (particularly daptomycin) in clinically relevant bacteria and the involvement of MprF in the virulence of pathogens, such as methicillin resistant *S. aureus*.

## Introduction

Gram-positive bacteria have a common cell envelope architecture that contributes to maintaining cell shape and protects the cell from environmental changes. The best characterised component of the cell envelope is the peptidoglycan (PG) which forms a mesh like structure enclosing the cell membrane. Onto this structure anionic polymers are either covalently attached to PG N-acetylmuramic acids and known as wall-teichoic acids (WTA, **Fig 1A**) or are tethered to the external face of the membrane and named lipoteichoic acids (LTA) (1–4). It is unclear whether LTA are localized in a periplasm-like space between the membrane and PG or extend through the PG layers (5, 6).

**Figure 1.**
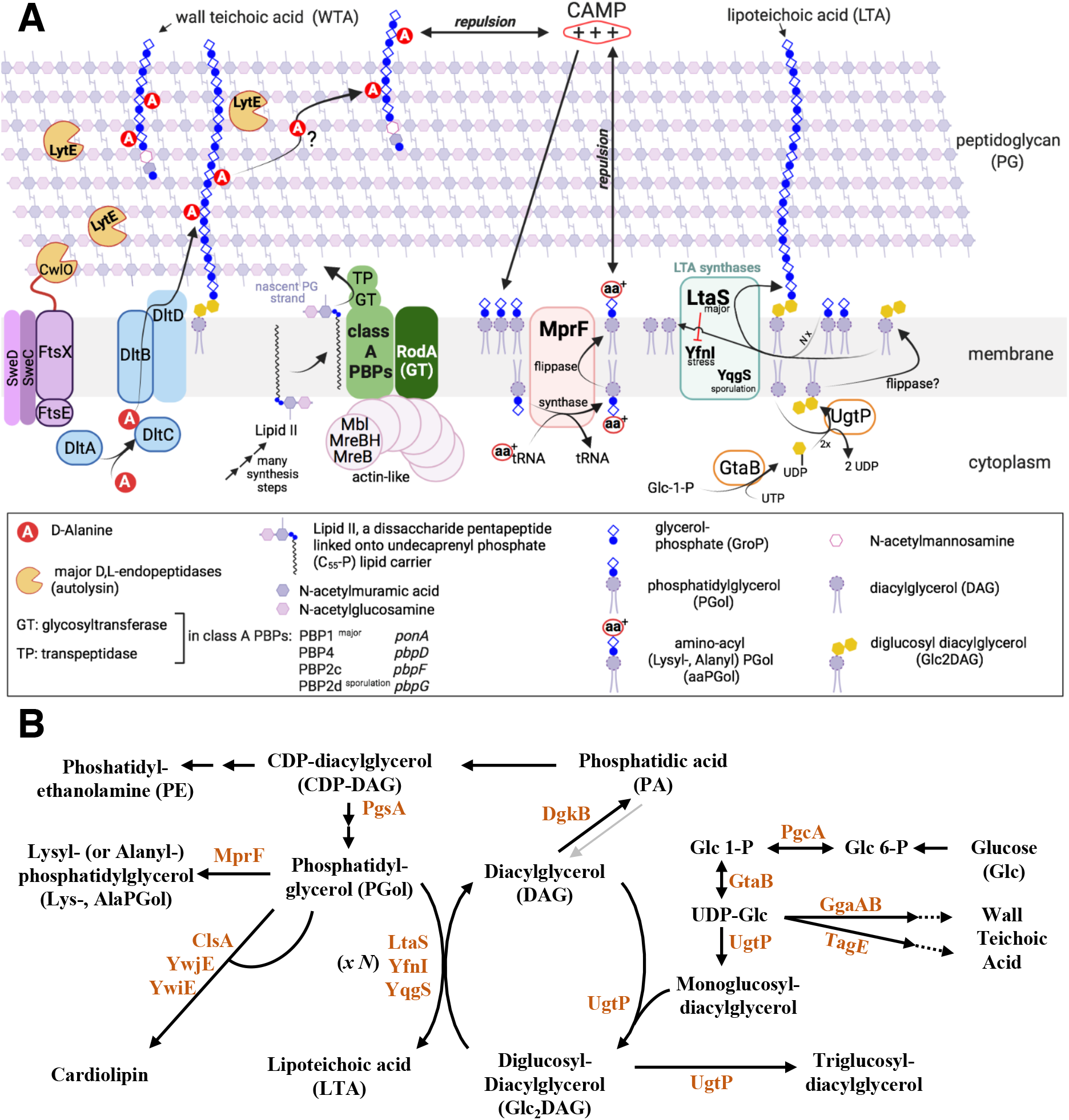
Representations of the cell envelope of Gram-positive rod-shaped bacterium *Bacillus subtilis* and the biosynthetic links between lipoteichoic acids and phospholipids. **(A)** Depiction of some of the cell envelope components and their function in *B. subtilis*. This figure indicates the complexity of the cell envelope synthesis due to the presence of enzymes that seemingly assume the same function. This redundancy of function has been identified in the reaction steps leading to the synthesis of the peptidoglycan (glycosyltransferases: 4 class A PBPs, PBP1/*ponA* major and the essential RodA), the lipoteichoic acids (3 LTA-synthases, where LtaS is the major enzyme) and the actin-like filaments that coordinate the cell shape (3 Mre-homologs: MreB/MreBH and Mbl where the latter is essential in *B. subtilis* 168CA). The major autolysins CwlO and LytE (the absence of both is lethal to *B. subtilis*) cleave the peptidoglycan pentapeptides. Presence of aminoacyl-phosphatidylglycerol (aaPGols) produced by MprF and the D-alanylation of teichoic acids by Dlt proteins are thought to create an electrostatic environment that protects the bacteria from CAMPs (cationic antimicrobial peptides). **(B)** A schematic representation of the biosynthetic links between lipoteichoic acid, phospholipids and glucolipids in *B. subtilis*, adapted from previous studies (3, 83). Lipoteichoic acid (LTA) consist of glycerolphosphate units anchored to the membrane by a glycolipid. In absence of cytoplasmic UgtP, LTA polymer is linked to the membrane by a diacylglycerol (DAG), instead of a diglucosyldiacylglycerol (Glc_2_DAG). When grown in LB at 37°C, the majority of *B. subtilis* phospholipids are phosphatidylglycerol (PGol) (37.5% ± 6.2), phosphatidylethanolamine PE (30.5% ± 3.8), Glc_2_DAG (9.6% ± 4.7) and lysyl-phosphatidylglycerol (LysPGol, 7.8% ± 3.7) produced by MprF (83). Recently, L- and N-succinyl-LysPGol, and also L- and D-AlaPGol were identified in *B. subtilis* (25). Grey arrow indicates a reaction step that might be bidirectional.

Peptidoglycan is synthetised by glycosyltransferases (GTases) which catalyse the addition of a disaccharide pentapeptide precursor (lipid II), to the nascent PG chain (**Fig 1A**) while transpeptidase (TPase) enzymes cross-link a proportion of the pentapeptides to adjacent PG strands (7, 8). In the absence of the bifunctional transmembrane GTase and TPase enzymes, known as class A Penicillin-Binding Proteins (aPBPs) (**Fig 1A**), *B. subtilis* cells are viable but the cells are longer and thinner and are prone to lysis (9, 10). The viability of *B. subtilis* lacking all aPBPs (referred as *ϕ..4*) depends on the functionality of the essential protein RodA that was recently identified to function as a PG glycosyltransferase (11, 12). The *ϕ..4* strain is conditionally lethal on PAB agar medium unless supplemented with magnesium (10). It is thought that this divalent cation alters the structure of the wall by binding to teichoic acids or PG and by doing so, might alter the activity of cell wall hydrolases (**Fig 1A**), the enzymes that modify and sculpt synthesised PG to allow the cell to elongate and divide (5, 13–15).

In rod-shaped *B. subtilis* 168, LTA and WTA are composed of polyglycerolphosphate (polyGroP) generated by independent biosynthetic pathways (**Fig 1B**) (2–5). Importantly, *B. subtilis* is not able to survive the loss of both LTA and WTA synthesis (16) and these teichoic acids seem to have roles in both cell growth (17) and division-separation (16, 18). It has also been determined that anionic charge of these polymers can be modified by the addition of D-alanine or N-acetylglucosamine (19). D-alanylation by Dlt proteins (**Fig 1A**) is proposed to regulate autolysin activity, cell wall ion homeostasis (3, 5) and also imparts some resistance to positively charged cationic antimicrobial peptides (CAMPs) (20). The net cell envelope charge is proposed to be moderated by the teichoic acids and also the phospholipid composition of the cell membrane (**Fig 1**) (21, 22). In this respect, the transmembrane protein MprF is the only enzyme known to synthesise and translocate aminoacyl-phosphatidylglycerol (aaPGol), in particular lysyl-PGol (**Fig 1**) (23–26). MprF plays a role in the virulence of bacterial pathogens (26, 27) and in *B. subtilis,* it confers resistance to CAMPs (**Fig 1A**) such as nisin and daptomycin through mechanisms that are not yet characterized (28, 29). Phosphatidylglycerol (PGol) is also the substrate for LTA-synthases, in *B. subtilis* three of these enzymes are present: LtaS (‘house-keeping’), YfnI (‘stress’) and YqgS (‘sporulation’) (16, 18) (**Fig 1**). These transmembrane LTA-synthases have an extracellular catalytic domain that uses PGol to generate a polyGroP polymer on a lipid anchor (2, 3). The lipid anchor is usually a diglucosyldiacylglycerol (Glc_2_DAG) produced by UgtP on the internal surface of the cell membrane (30) (**Fig 1A**). As an *ugtP* mutant still produces LTA (31), it suggests that polyGroP polymers are probably anchored to a phosphatidylglycerol as observed in *Staphylococcus aureus* (32, 33).

In *S. aureus,* LTA is essential and seems to modulate autolytic activity (34), while WTA is dispensable but appears to restrict autolytic activity (1, 4). Contrary to *B. subtilis*, *S. aureus* has only one LTA-synthase (LtaS) whose absence is (conditionally) essential (35) and can be complemented by *B. subtilis ltaS* but not *yfnI* (18, 36). A recent *in vitro* study showed that *S. aureus* LtaS is processive and that the LTA polymer elongation is directly regulated by the identity and concentration of lipid starter units. Interestingly, use of Glc_2_DAG as a starter unit led to the formation of shorter polymers (33) while the use of PGol resulted in the synthesis of longer polymers. It is not clear if this newly identified mechanism might be conserved in *B. subtilis.* However, in absence of *B. subtilis* LtaS, YfnI activity results in LTA of increased length and it is possible that the combination of LtaS and YfnI activities modulated the length of LTA (**Fig 1A**) (18, 37).

Through characterising *B. subtilis* strains lacking peptidoglycan GTase activity (12), we aimed to understand the cell wall stresses causing aPBP mutants to lyse on glucose rich media. Collectively, our data supports the idea that in *B. subtilis* LTA length acts to moderate the activity of at least one important autolysin, LytE (**Fig 1A**). The presence of glucose in a strain lacking aPBPs increases the expression of LtaS and YfnI and these result in altered LTA production. Crucially, we identify a new role for MprF in altering LTA polymerisation in *B. subtilis* and *S. aureus*. Our results reveal that MprF acts to change the cell envelope in a more dramatic way than simply increasing the positive charge on the cell membrane which has implications for understanding the association of MprF with both virulence and antibiotic resistance in Gram-positive bacteria.

## Results

### MprF alters cell viability in absence of *ponA* or *mbl*

In *Bacillus subtilis*, the absence of all aPBPs (*τ1.4* for *τ1.ponA τ1.pbpD τ1.pbpF τ1.pbpG* mutant) is conditionally lethal on glucose-rich PAB agar medium unless supplemented with magnesium (10, 12, 38). The suppressive effect of divalent magnesium has been observed in other cell-wall deficient mutants (10, 16, 38, 39) and it is not well understood. To identify the cell wall factors able to suppress *τ1.4* lethality, we applied a random transposition mutagenesis to a strain deleted for the vegetative aPBPs (**Tables 1-3**). Mapping of the transposon in strain SWA11a revealed an insertion in the gene *mprF*, encoding a lysyl-phosphatidylglycerol (LysPGol) synthase known to participate in phospholipid metabolism (**Fig 1**) (23, 28). Introduction of a *mprF* deletion into *τ1.4*, confirmed that *τ1.4 τ1.mprF* was viable on nutrient agar (NA) with glucose and on PAB agar (**Fig 2A, Fig S1A**). A strain lacking *ponA* (encoding the major aPBP (PBP1) is viable but exhibits a growth phenotype that is suppressed by magnesium (**Fig 2A, Fig S1B**). Here the deletion of *mprF* in *τ1.ponA* improved its growth and its colony morphology.

**Figure 2.**
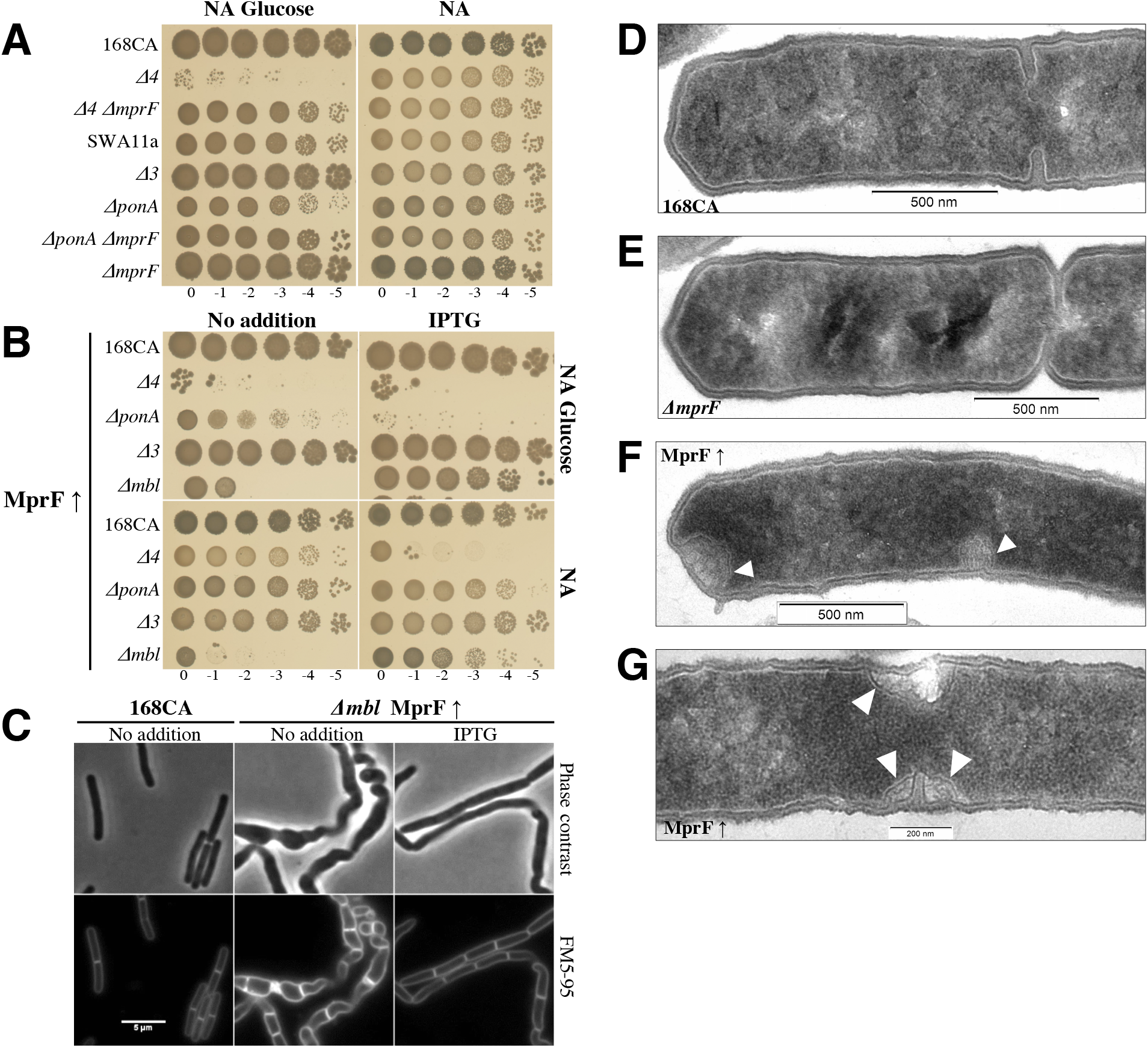
MprF alters cell viability of *B. subtilis* class A PBPs and *mbl* mutants. **(A)** Ten-fold spot growth assays of strains *B. subtilis* wild type 168CA, AG157 (*ΔponA ΔpbpD ΔpbpF ΔpbpG* renamed *Δ4*), AG223 (*Δ4 ΔmprF*), SWA11a (*ΔponA ΔpbpD ΔpbpF mprF::TnYLB-1 lacA*), AG417 (*ΔpbpD ΔpbpF ΔpbpG* renamed *Δ3*), RE101 (*ΔponA*), AG193 (*ΔponA ΔmprF)* and AG181 (*ΔmprF*). Plates (here glucose 1%) were incubated at 37°C for 24 h and images taken. **(B)** The effect of *mprF* overexpression (MprF t) is assay in different strain backgrounds using an IPTG-inducible *mprF* construct. Plates (here glucose 1%) were incubated at 37°C for 24 h and scanned. Strains tested were AG304 (168CA MprF t), AG317 (*Δ4* MprF t), AG311 (*ΔponA* MprF t), AG421 (*Δ3* MprF t) and AG322 (*Δmbl* MprF t). **(C)** Phase-contrast and membrane dye (FM5-95) microscopy images of *B. subtilis* 168CA and AG322 (*Δmbl* MprF t) strains grown in NB ± IPTG (0.1 mM) to mid-exponential phase at 37°C. Scale bar indicates 5 µm. **(D-G)** Transmission electron microscopy (TEM) images show cross-sections of (**D**) wild type 168CA, (**E**) AG181 (*ΔmprF)* and (**F-G**) AG304 strain (MprF t) grown in nutrient broth (and IPTG for AG304). Arrows indicate abnormal cell wall structures that seem specific to the strain MprF t. All images were taken at 15,000x magnification and processed in Fiji with scale bars attached. Data information. Strains phenotype was tested at least three times, TEM experiment was carried out one time. Supporting information for this figure is presented in **Fig S1, S2AB**.

**TABLE 1.** Plasmids used in this study (by alphabetic order)

| Plasmids | Relevant features<sup>a</sup> | Source or reference |
| --- | --- | --- |
| pAM-21 | *kan* *P*<sub>T7</sub> *LytE*<sup>(26-334AA)</sup>-6His *lacI* | This study |
| pCotC-GFP | used as matrix to amplify *cat* | (75) |
| pDR111 | *bla* *amyE*::(*lacI* *P*<sub>hyspank</sub> *spc*) | Rudner, D., unpublished |
| pDR111-*P*<sub>hyspank</sub>-*ltaS* | *bla* *amyE*::(*lacI* *P*<sub>hyspank</sub>-*ltaS*, *spc*) | This study |
| pDR111-*P*<sub>hyspank</sub>-*mprF* | *bla* *amyE*::(*lacI* *P*<sub>hyspank</sub>-*ATGmprF*, *spc*) | This study |
| pDR111-*P*<sub>hyspank</sub>-*yfnI* | *bla* *amyE*::(*lacI* *P*<sub>hyspank</sub>-*yfnI*, *spc*) | This study |
| pDR111-*P*<sub>hyspank</sub>-*yqgS* | *bla* *amyE*::(*lacI* *P*<sub>hyspank</sub>-*yqgS*, *spc*) | This study |
| pET28a (+) | *kan* *P*<sub>T7</sub> *his* *lacI* | Novagen |
| pG<sup>+</sup>host9::*ΔponA* | *erm*, carry flanking DNA regions of *ponA* | (12) |
| pG<sup>+</sup>host10::*ΔpbpD* | *bla*, carry flanking DNA regions of *pbpD* | (12) |
| pG<sup>+</sup>host10::*ΔpbpF* | *bla*, carry flanking DNA regions of *pbpF* | (12) |
| pIC22 | used as matrix to amplify *phleo* | (76) |
| pLOSS* | *bla* *spc* *P*<sub>spac</sub>-*mcs* *P*<sub>divIVA</sub>-*lacZ* *lacI*<br>*rep*<sub>pLS20</sub>(*GA*→*CC*) | (39) |
| pLOSS-*P*<sub>spac</sub>-*lytE* | *bla* *spc* *P*<sub>spac</sub>-*lytE* *P*<sub>divIVA</sub>-*lacZ* *lacI*<br>*rep*<sub>pLS20</sub>(*GA*→*CC*) | This study |
| pLOSS-*P*<sub>spac</sub>-*ponA* | *bla* *spc* *P*<sub>spac</sub>-*ponA* *P*<sub>divIVA</sub>-*lacZ* *lacI*<br>*rep*<sub>pLS20</sub>(*GA*→*CC*) | (12) |
| pMarB | *bla* *erm* TnYLB-1::*kan* *HimarI* transposase | (77) |
| pMUTinHis | *bla* *erm* *P*<sub>spac</sub> *mcs* 12x*his* *lacZ* *lacI* | (73) |
| pMUTin-*lytE*-*his* | *bla* *erm* *P*<sub>spac</sub> *mcs* *lytE*-6AA-12His *lacZ* *lacI* | This study |
| pMUTin-*ltaS*-*his* | *bla* *erm* *P*<sub>spac</sub> *mcs* *LtaS*-6AA-12His *lacZ* *lacI* | This study |
| pMUTin-*yfnI*-*his* | *bla* *erm* *P*<sub>spac</sub> *mcs* *YfnI*-6AA-12His *lacZ* *lacI* | This study |

<sup>a</sup>*cat*, chloramphenicol resistance; *bla*, ampicillin resistance; *spc*, spectinomycin resistance; *erm*, erythromycin resistance; *phleo*, phleomycin resistance; *kan*, kanamycin resistance; *lacI*, encodes lactose repressor; *mcs*, multiple cloning site; *lacZ*, encodes β-galactosidase; AA, amino acids; *P*<sub>hyspank</sub> and *P*<sub>spac</sub> are IPTG-inducible promoters with *P*<sub>hyspank</sub> known to be stronger; *rep*<sub>pLS20</sub>(*GA*→*CC*), unstable replication in *Bacillus*.

In the light of these results we reasoned that if MprF activity is detrimental to *τ1.4* its overexpression should be deleterious to *τ1.ponA*. Consistent with this, the overexpression of MprF (MprF t) was found to be lethal in *ponA-*deleted strains on NA-glucose (**Fig 2B**). In contrast, on NA, both *τ1.ponA* and *τ1.4* strains grew slowly when *mprF* overexpression was induced (**Fig 2B**) but no obvious effect was observed for 168CA or *Δ3* aPBPs (*ΔpbpD ΔpbpF ΔpbpG*). We also observed that the overexpression of *mprF* caused the cells of *ΔponA* and *Δ4* to lyse in nutrient broth (NB) (**Fig S1C**).

The identification of *ΔmprF* as a suppressor suggested that increased negative charge on the membrane could contribute to the rescue of *Δ4* growth. This seemed to contradict the long-standing hypothetical role of magnesium in altering the cell envelope by neutralizing anionic polymers and/or autolysis activity (1, 5). As it is known that the function of *dlt* operon is to add positive charge to LTA, it would be predicted that the loss of Dlt’s function should have a similar effect as *ΔmprF* on *Δ4* strain viability (40, 41). However, *Δ4 ΔdltAB* strain was unable to grow on glucose rich media (data not shown). We also deleted the genes encoding the cardiolipin synthases in *Δ4* to alter the membrane charge and the cellular pool of PGol (**Fig 1B**) but the loss of cardiolipin had no suppressive effect on the *Δ4* phenotype (data not shown).

To understand if the effect of MprF was specific to *Δ4*, we altered MprF expression in other known magnesium dependent mutants *ΔmreB* and *Δmbl*. It should however be noted that only *Δmbl* requires magnesium to grow on plate compared to *ΔmreB* (**Fig S2A**). MreB, Mbl and MreBH are actin-like isologues which help maintain cell shape by controlling both cell wall synthesis (42) and the major autolytic enzymes CwlO and LytE (Mbl for CwlO and MreB/BH for LytE) (43). Interestingly, we identified that MprF overexpression rescued the growth of *Δmbl* on NA-glucose and NA (**Fig 2B**). In addition, the morphological defects associated with this mutation (16) were less pronounced, resulting in wide rod-like shape cells and cell chains that were less often twisted (**Fig 2C**).

To explain our genetic findings in *Δmbl* background, we had to consider that MprF might play a role in cell wall metabolism. This idea seemed to be supported by transmission electron microscopy (TEM) of a strain overexpressing MprF (**Fig 2FG**). Here cells exhibited an increase in cell wall thickness (it was also observed that cell poles were ‘dented’, and septa were misplaced), a defect similar to that observed in cell division mutants. However the change in cell morphology of the strain deleted for *mprF* was less significant (**Fig 2E**).

Previous reports showed that lethality of *B. subtilis* Δ*mbl* is suppressed by the deletion of the major LTA-synthase LtaS (16) and other work had shown that in *ltaS* mutant the activity of the LTA-synthase YfnI results in longer LTA polymers (18). This suggested that the overexpression of MprF in Δ*mbl* might diminish the cellular level of PGol (through a different pathway, **Fig 1B**) (24) or MprF might function in LTA synthesis.

### Conditional essentiality of UgtP in the absence of MprF and the Class A PBPs

To comprehend the effect of MprF-loss in a strain lacking aPBPs and identify the MprF suppression pathway, we carried out a transposon screen in a strain lacking the vegetative class A PBPs, *mprF, lacA* and carrying pLOSS-*P_spac_*-*ponA* (strain AG200, **Tables 1-2**) (12) with the objective of identifying alleles that rendered the maintenance of the plasmid copy of *ponA* essential. From this screen strain AG200BK#42 was isolated with the expected phenotype and found to carry a transposon inserted in the gene *gtaB*. GtaB catalyses a cytoplasmic conversion of Glucose-1-P to UDP-Glc (**Fig 1**), where the latter ultimately contributes to LTA synthesis and WTA modification. Subsequent mutagenesis in backgrounds *Δ4* and *Δ4 ΔmprF* confirmed the lethal (NA-glucose) and sick (NA) phenotypes in absence of GtaB (**Fig 3A**).

**Figure 3.**
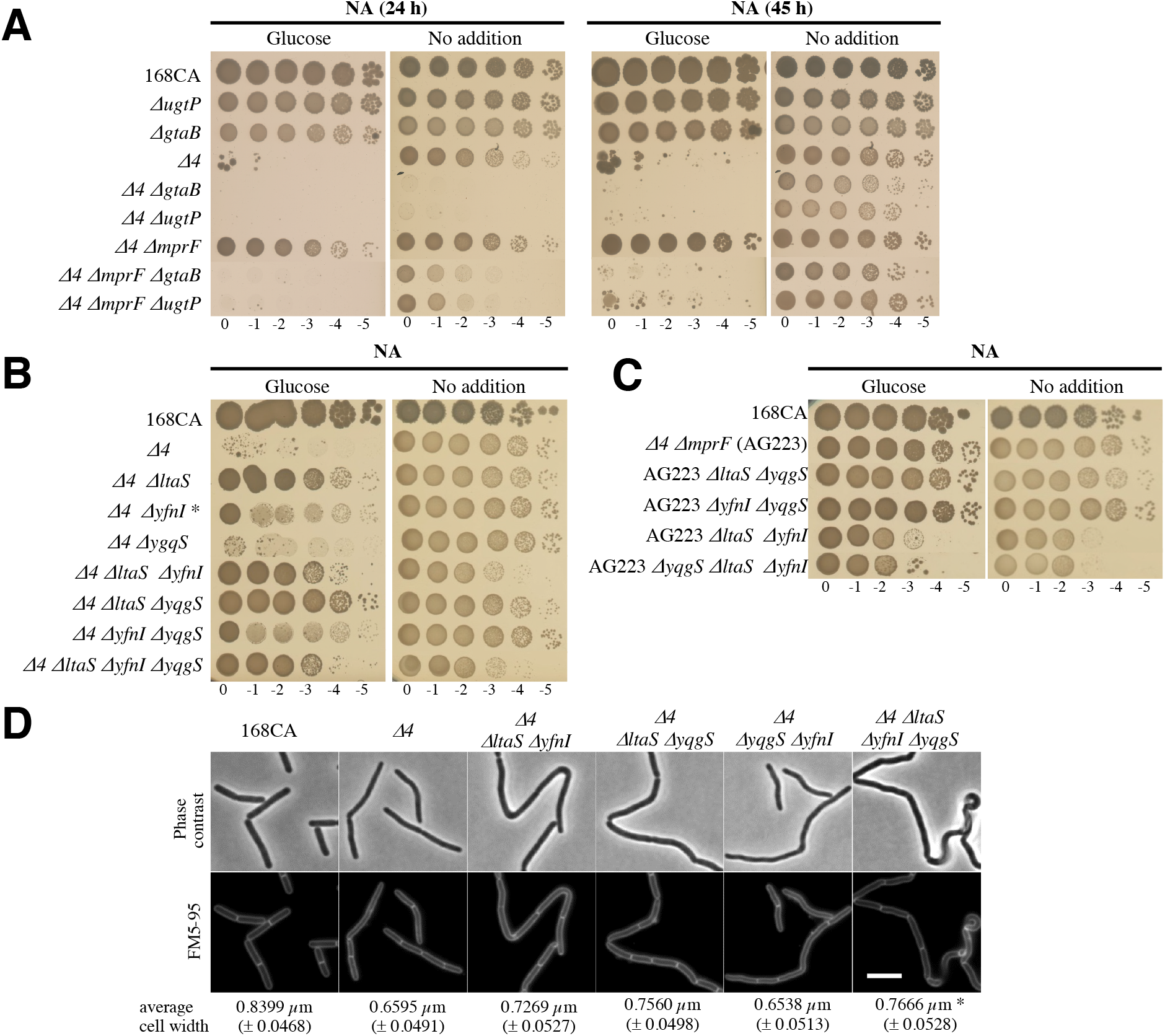
LtaS causes cell morphogenesis defect of an all-class A PBP mutant. **(A)** Absence of *gtaB* or *ugtP* causes severe growth defect in absence of class A PBPs and MprF. Strains were grown exponentially in NB with 20 mM MgS0_4_, then were washed and diluted in NB prior to preparing a 10-fold serial dilution in NB spotting on the NA and NA glucose (0.5%) plates which were incubated at 37°C and imaged after 24 h and 45 h. Strains were also spotted on same media supplemented with 10 mM MgS0_4_ to confirm all the strains dilution spots could grow, at 17 h of incubation only *Δ4 ΔgtaB* and *Δ4 ΔugtP* displayed colonies of smaller size at that time point (data not shown). Strains tested: *B. subtilis* wild type (168CA), PG253 (*ΔugtP*), SM08 (*ΔgtaB*), AG157 (*Δ4*), AG290NopLOSS (*Δ4 ΔgtaB*), AG443 (*Δ4 ΔugtP*), AG223 (*Δ4 ΔmprF*), AG636 (*Δ4 ΔmprF ΔgtaB*) and AG632 (*Δ4 ΔmprF ΔugtP*). **(B)** Spot dilution growth assays for the following strains: *B. subtilis* wild type 168CA, AG157 (*Δ4*), AG342 (*Δ4 ΔltaS*), AG343* (*Δ4 ΔyfnI*), AG344 (*Δ4 ΔyqgS*), AG370 (*Δ4 ΔltaS ΔyfnI*), AG372 (*Δ4 ΔltaS ΔyqgS*), AG377 (*Δ4 ΔyfnI ΔyqgS*) and AG380 (*Δ4 ΔltaS ΔyfnI ΔyqgS*). Samples were inoculated onto NA ± glucose 1% and, incubated at 37°C for 24 h. * This strain was found to pick up suppressor mutations more rapidly than AG342. **(C)** Ten-fold serial dilutions were prepared for *B. subtilis* wild type 168CA, AG223 (*Δ4 ΔmprF*), AG389 (AG223 *ΔltaS ΔyqgS*), AG390 (AG223 *ΔyfnI ΔyqgS*), AG399 (AG223 *ΔltaS ΔyfnI*) and AG400 (AG223 *ΔyqgS ΔltaS ΔyfnI*) strains. Plates (here glucose 1%) were incubated at 37°C and imaged after 24h. **(D)** Representative microscopy images of cells of strains of (A) grown in NB at 37°C for 120 min and stained with a FM5-95 dye. Scale bar represents 5 µm. The average of cells diameter extracted from **Table 4** is indicated. * The cells measurement was done on the straight regions of the cell chain. Data information: Strains phenotype was tested at least three times. Supporting information is presented in **Fig S2C, S3** which also display results obtained in *ΔponA* background.

**TABLE 4.**
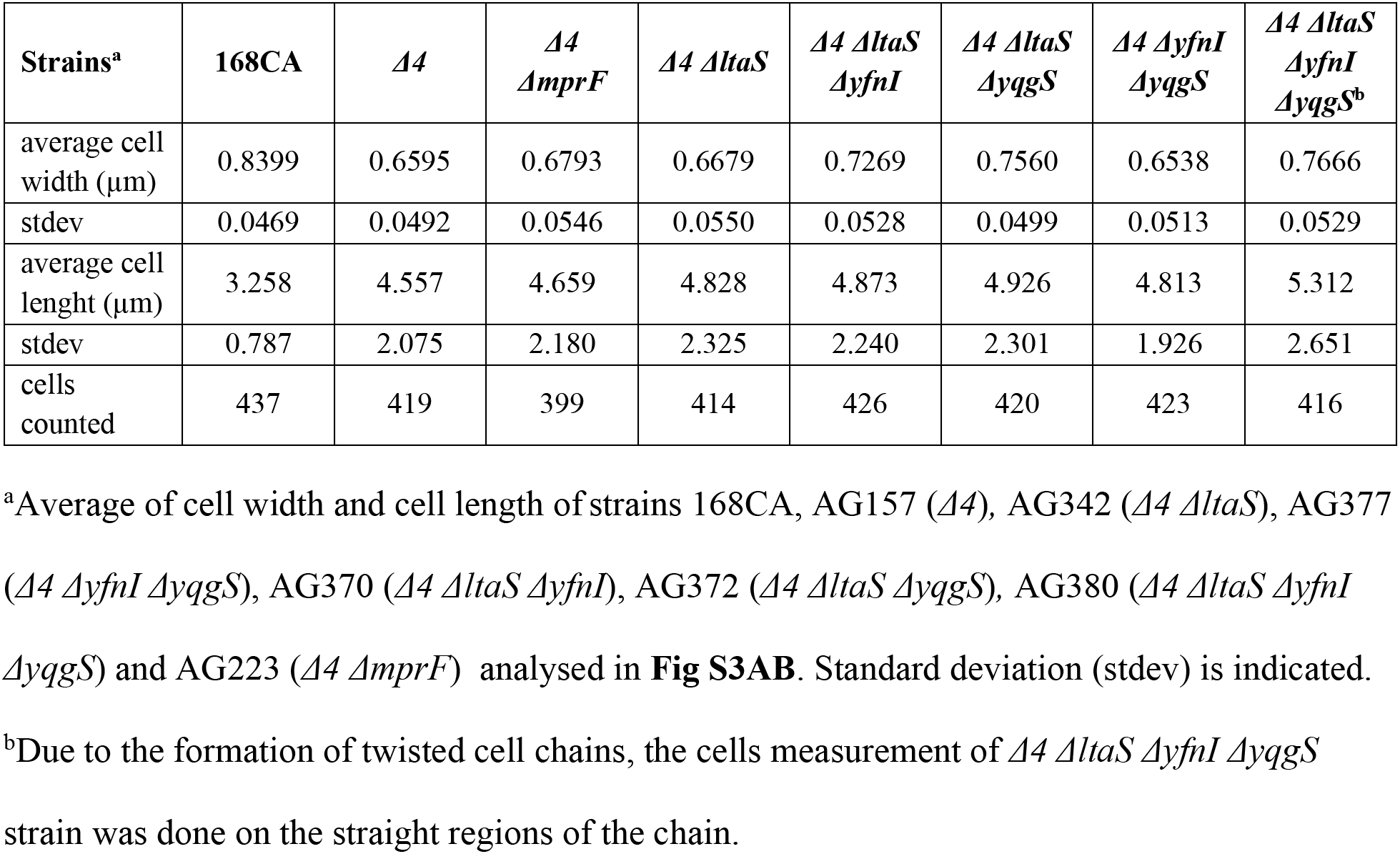
Cell morphogenesis changes associated with deletion of LTA-synthase genes combined with class A PBPs null mutants of *B. subtilis*

| Strains<sup>a</sup> | 168CA | $\Delta 4$ | $\Delta 4$<br>*AmprF* | $\Delta 4$ ΔltaS | $\Delta 4$ ΔltaS<br>ΔyfnI | $\Delta 4$ ΔltaS<br>ΔyqgS | $\Delta 4$ ΔyfnI<br>ΔyqgS | $\Delta 4$ ΔltaS<br>ΔyfnI<br>ΔyqgS<sup>b</sup> |
| --- | --- | --- | --- | --- | --- | --- | --- | --- |
| average cell width (μm) | 0.8399 | 0.6595 | 0.6793 | 0.6679 | 0.7269 | 0.7560 | 0.6538 | 0.7666 |
| stdev | 0.0469 | 0.0492 | 0.0546 | 0.0550 | 0.0528 | 0.0499 | 0.0513 | 0.0529 |
| average cell length (μm) | 3.258 | 4.557 | 4.659 | 4.828 | 4.873 | 4.926 | 4.813 | 5.312 |
| stdev | 0.787 | 2.075 | 2.180 | 2.325 | 2.240 | 2.301 | 1.926 | 2.651 |
| cells counted | 437 | 419 | 399 | 414 | 426 | 420 | 423 | 416 |

<sup>a</sup>Average of cell width and cell length of strains 168CA, AG157 ( $\Delta 4$ ), AG342 ( $\Delta 4$ ΔltaS), AG377 ( $\Delta 4$ ΔyfnI ΔyqgS), AG370 ( $\Delta 4$ ΔltaS ΔyfnI), AG372 ( $\Delta 4$ ΔltaS ΔyqgS), AG380 ( $\Delta 4$ ΔltaS ΔyfnI ΔyqgS) and AG223 ( $\Delta 4$ *AmprF*) analysed in **Fig S3AB**. Standard deviation (stdev) is indicated.
<sup>b</sup>Due to the formation of twisted cell chains, the cells measurement of $\Delta 4$ ΔltaS ΔyfnI ΔyqgS strain was done on the straight regions of the chain.

To determine if this was related to LTA synthesis or WTA modification, we deleted sequentially, in the *Δ4* or *Δ4* pLOSS-*P_spac_*-*ponA* backgrounds, each enzyme whose function is immediately upstream or downstream of GtaB (**Fig 1B)**. Characterisation of the resulting strains (**Table 2**) showed that only deletion of *ugtP*, the enzyme which produces the Glc_2_DAG lipid anchor for LTA, had a similar effect to that of *gtaB*. Strain *Δ4 ΔmprF ΔugtP* was unable to grow on NA glucose (**Fig 3A**), a phenotype common to *ΔponA ΔgtaB* and *ΔponA ΔugtP* (**Fig S2C**). On NA, *Δ4 ΔgtaB* and *Δ4 ΔugtP* grew very slowly (**Fig 3A,** right panel).

**TABLE 2.**
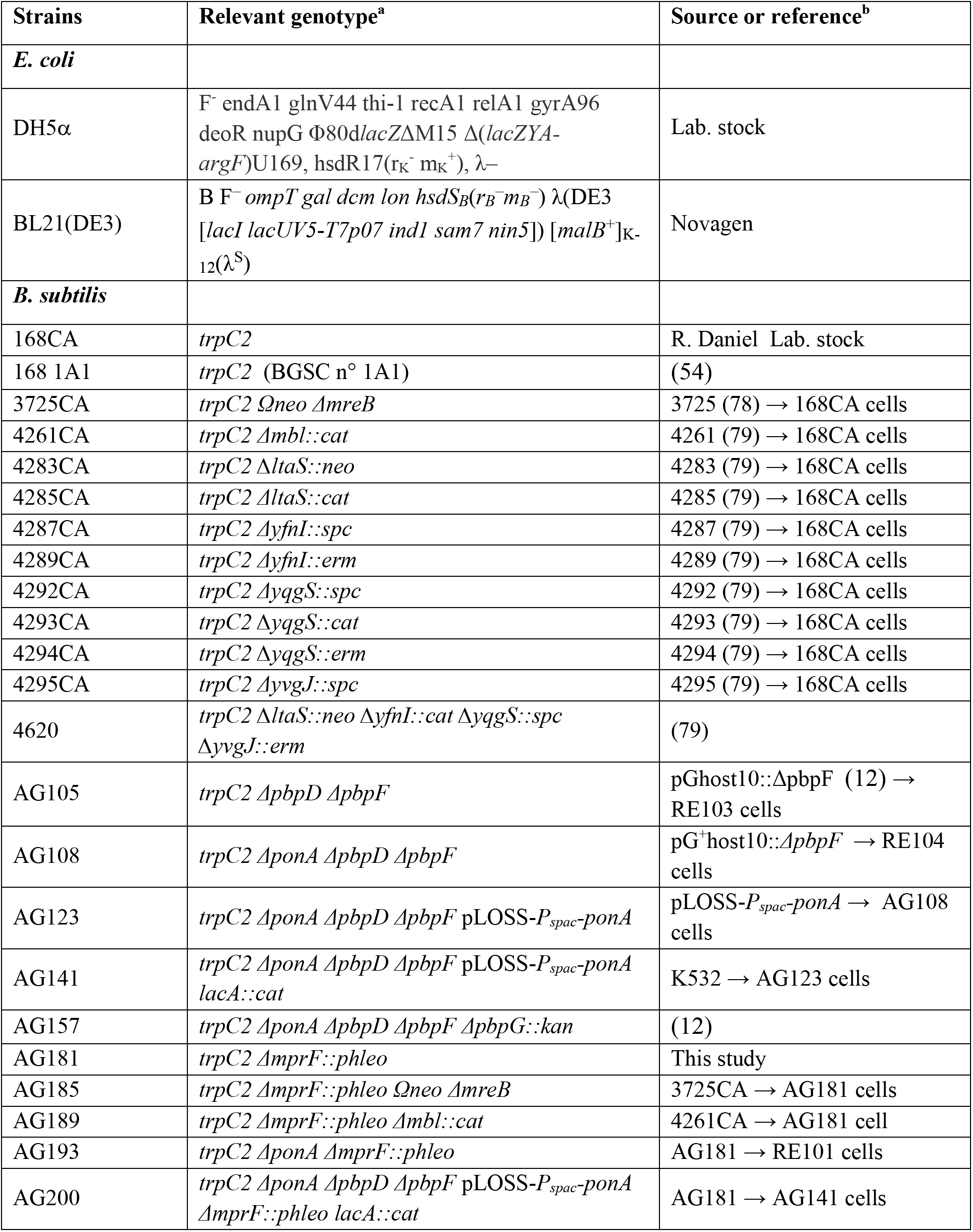
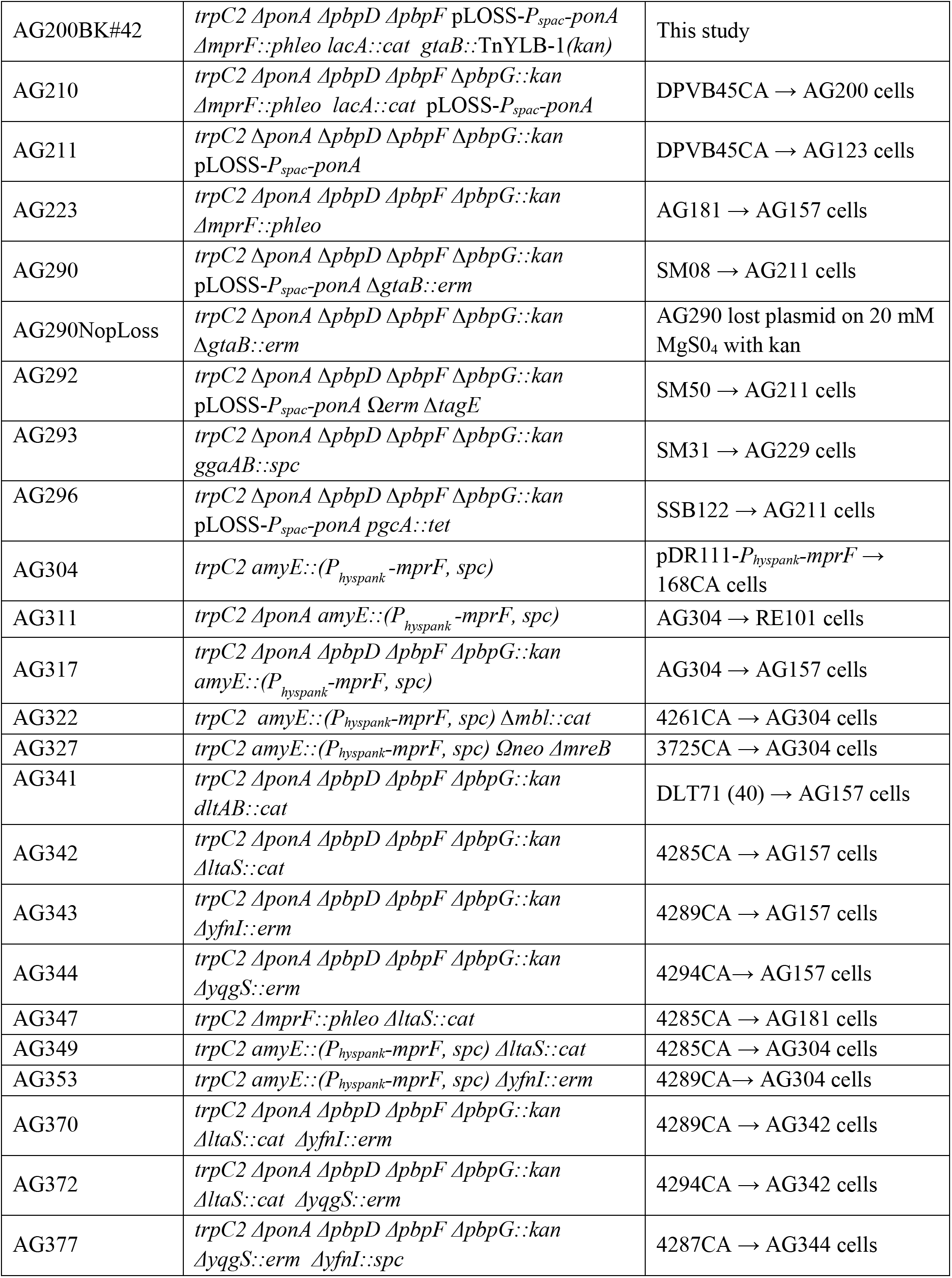
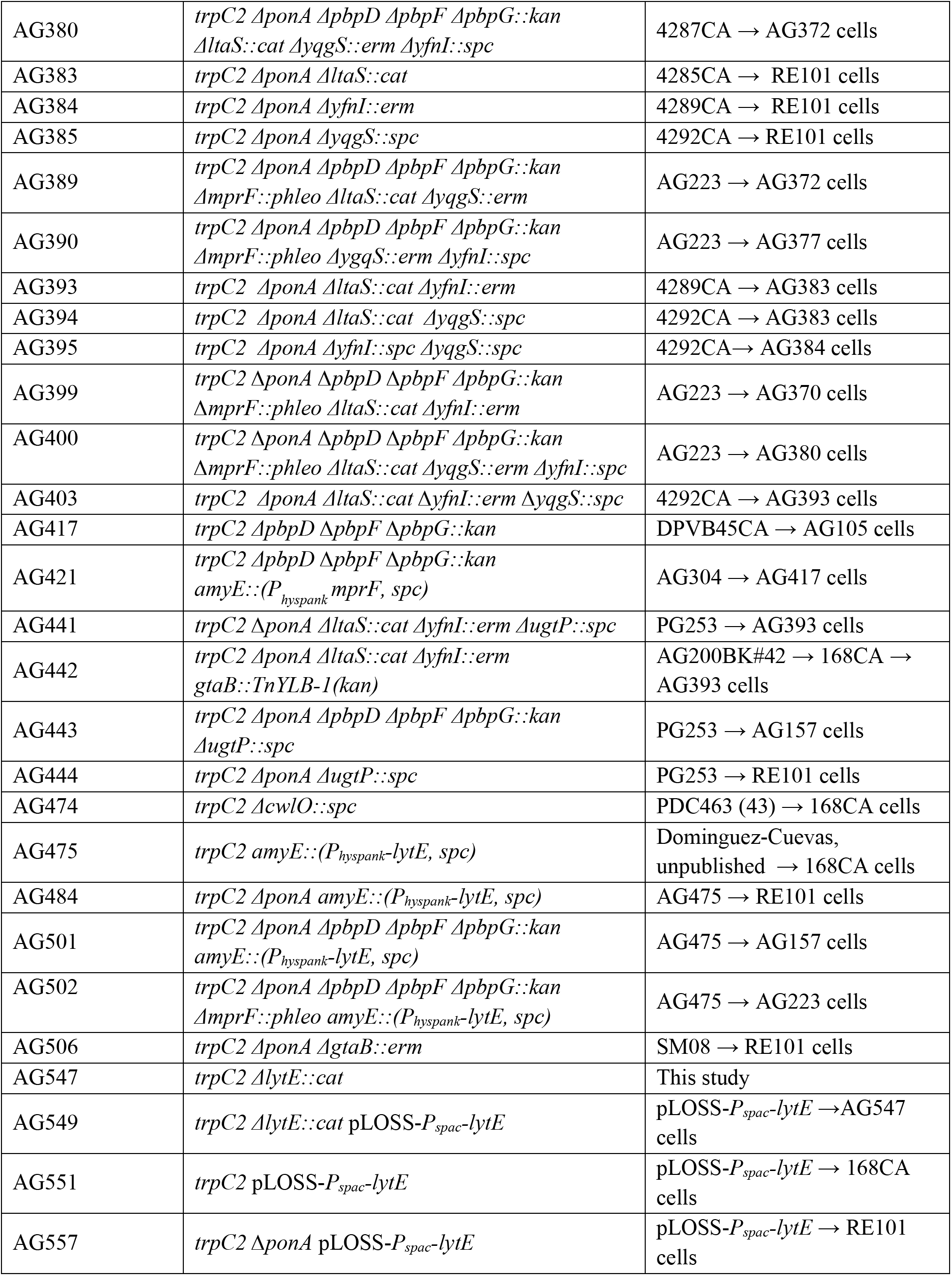
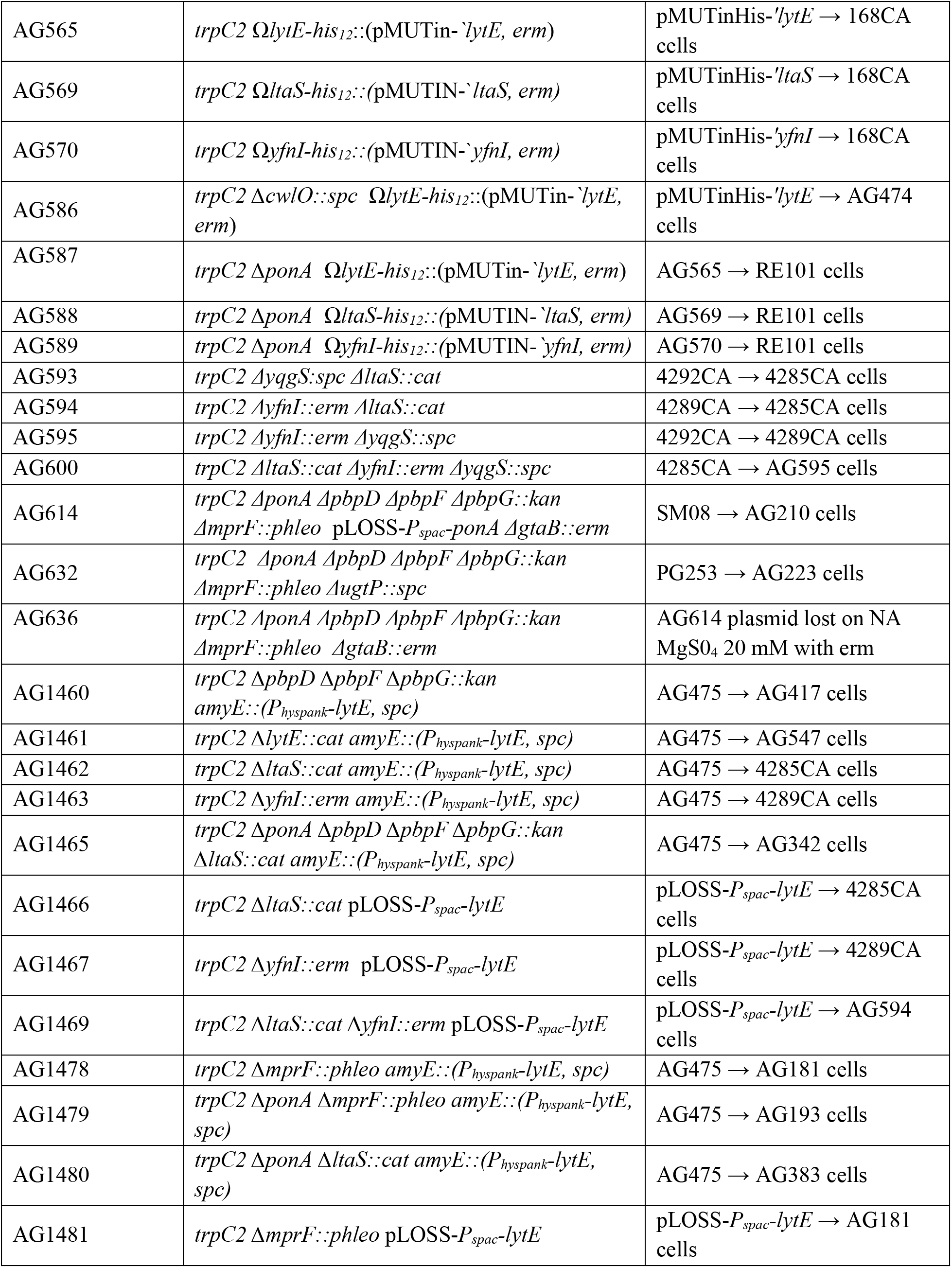
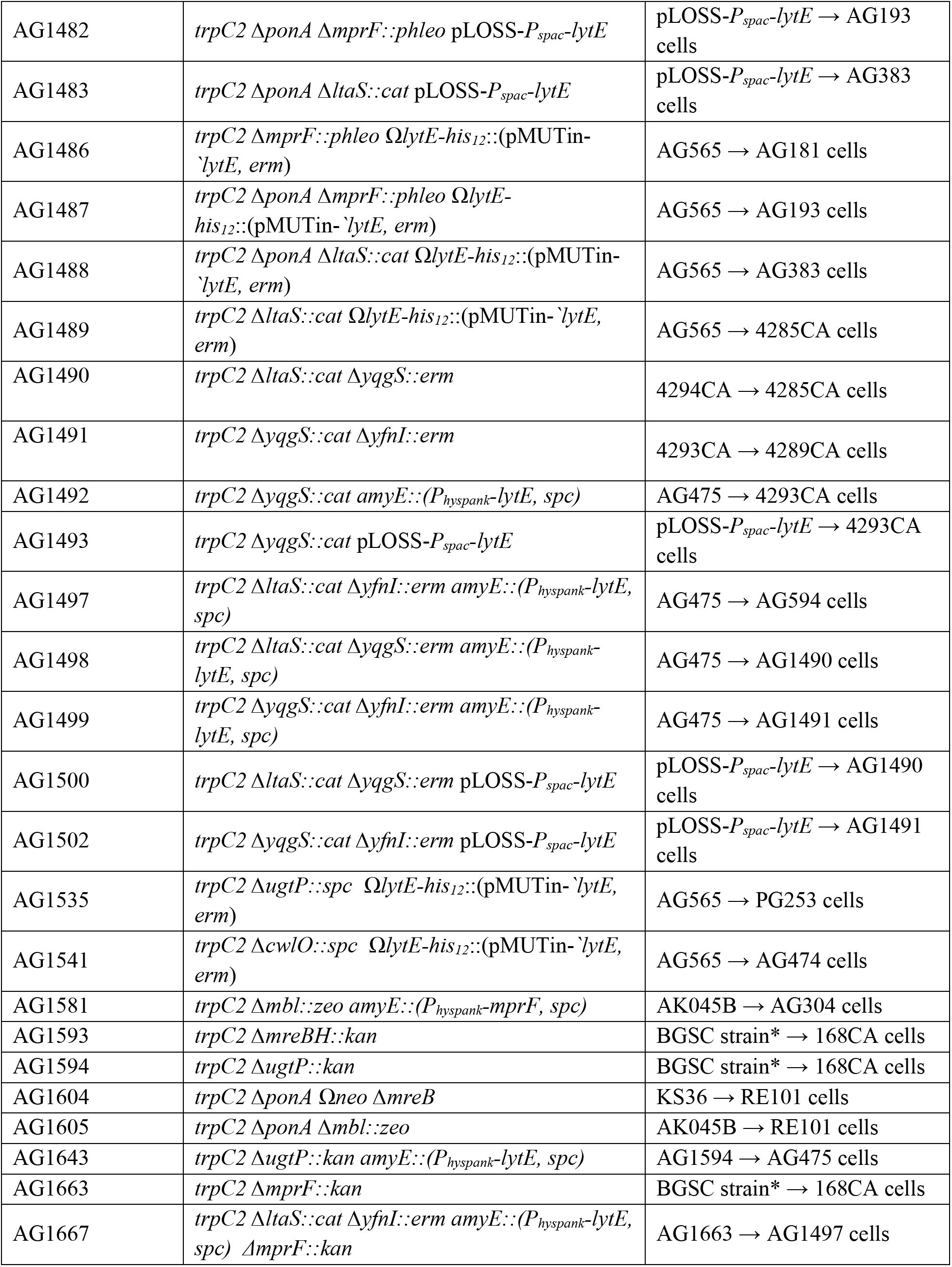
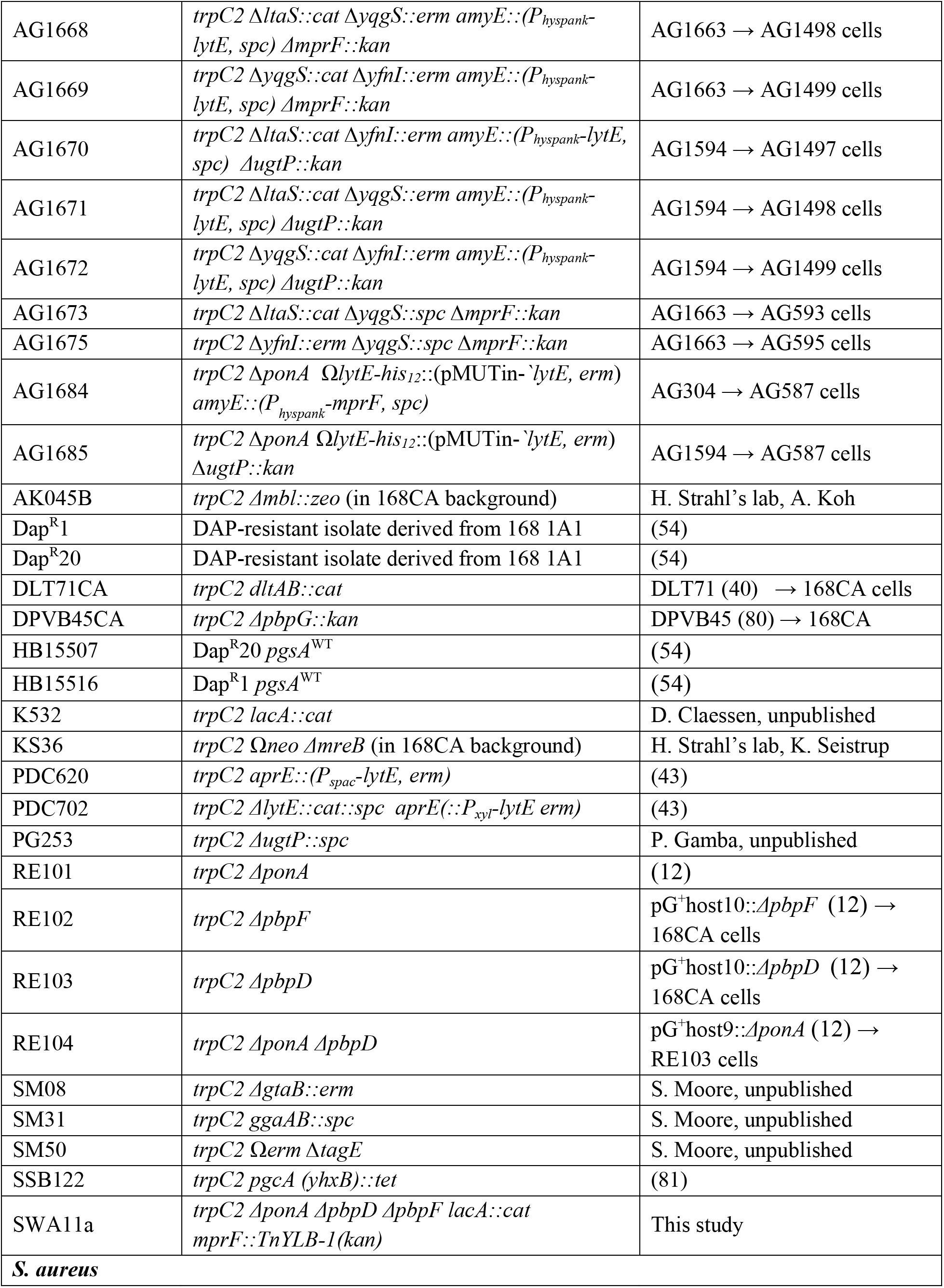
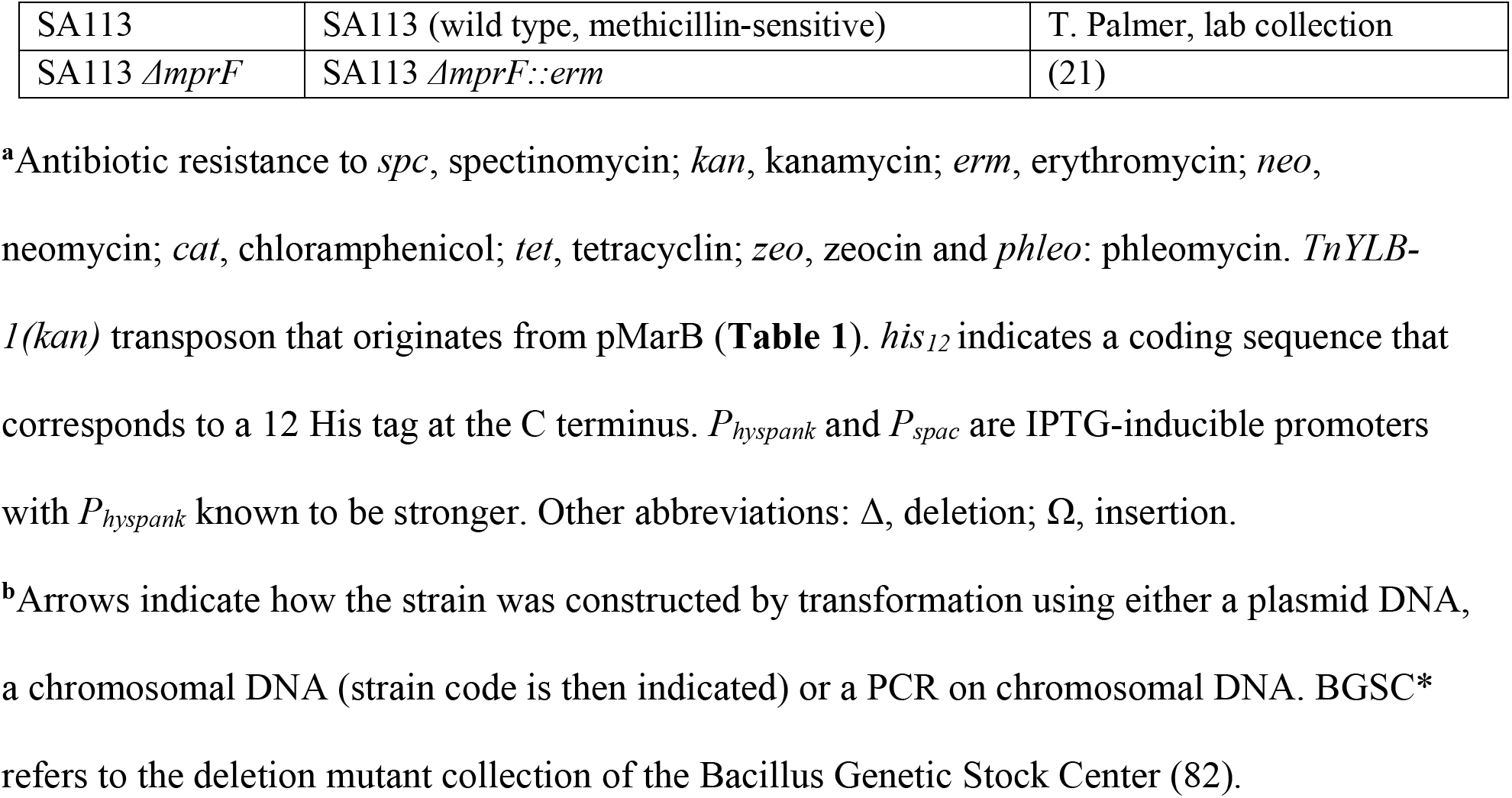
Strains used in this study

LysPGol content increases in *ΔugtP* cells (24) which could explain the similarity of phenotype of *Δ4 ΔugtP* and *Δ4* MprF 1’ (i.e. slow growing on NA) but not the dominant effect of *ΔugtP* on *Δ4 ΔmprF.* In *S. aureus,* it has been observed that *ΔugtP* (*ypfP*) and *ΔgtaB* mutants produce long LTA (32). Thus this context and our results reinforced the idea that MprF may have a role in LTA synthesis.

### Growth of a strain lacking all aPBPs mutant improves in absence of LtaS

To verify that LTA synthesis alters *Δ4* growth, we analysed the phenotype of strains lacking aPBPs and the LTA-synthase genes individually or in combination (**Fig 1**). Surprisingly, the *Δ4 ΔltaS* mutant was viable on NA-glucose as well as with the additional deletion of *yfnI* or *yqgS* (**Fig 3B**). Other strains expressing LtaS showed limited growth and were found to pick up suppressor mutations rapidly on glucose (e.g. *Δ4 ΔyfnI*, *Δ4 ΔyfnI ΔyqgS,* **Fig 3B**). Thus, LtaS activity is also linked to the glucose-associated lethality of *11.4.* Interestingly, the absence of MprF is more advantageous to *Δ4,* as it results in a faster growth and larger colonies formed on NA-glucose compared to *Δ4 ΔltaS* (**Fig S1B**). This phenotypic difference indicated that slightly different mechanisms helped restore the growth of *11.4*.

To test this, LTA-synthase genes were systematically deleted in the *Δ4 ΔmprF* (AG223) background and the viability of the strains generated determined on NA with or without glucose. All the strains remained viable on NA glucose (**Fig 3C**) which indicated the effect of MprF loss could be related to LTA. In support of this, in the *Δ4 ΔmprF* background the combined absence of *ltaS* and *yfnI* delayed growth on NA glucose resulting in a phenotype similar to that observed on NA (**Fig 3C)**. Thus, in these specific genetic backgrounds LTA synthesis is not essential but permitted higher growth efficiency.

Neither the absence of LtaS nor MprF had any effect on the filamentous thin cell morphology of *Δ4* (**Fig S3A-B, Table 4**) but cell lysis was significantly decreased in NB. In the absence of all LTA-synthases, *Bacillus* is viable but the strain forms filamentous and clumpy cells (16). We observed that the combined absence of LtaS, YfnI and/or YqgS in *Δ4, ΔponA* or *Δ4 ΔmprF* genetic backgrounds lead to an increase in cell diameter and cells remained linked together at the site of division (**Fig 3D, Fig S3A-B, Table 4**). However, when LtaS was the only functional LTA-synthase (e.g. *Δ4 ΔyqgS ΔyfnI*, **Fig 3D**), cells were thin and comparable to *Δ4* and the *ΔponA* background (**Fig S3C-D**). Collectively, our results indicate that the activity of the LTA synthases, in particular LtaS activity, could account for *Δ4* phenotype.

### Impact of glucose, PBP1 and MprF on LTA synthesis

To comprehend the effect of LtaS, MprF and UgtP in the absence of aPBPs we decided to analyse LTA production. We developed a Western blot protocol relying on a monoclonal antibody for Gram-positive LTA (Thermofischer) and a polyclonal PBP2B antibody. First, we determined the LTA profile for 168CA and the LTA-synthases mutants (**Fig 4A** and **Fig S4**) grown in NB and NB glucose (0.2%). The wild type LTA signal corresponded to a mobility comparable to 7-17 kDa as previously identified (18). Interestingly, our media comparison showed that LTA-signal in *ΔltaS* cells grown in NB-glucose (> 15 kDa) (**Fig 4A** left panel, 2^nd^ lane) was distinct from that seen for growth in NB (7-32 kDa) (**Fig 4A** right panel, 2^nd^ lane). However, the LTA pattern was different when YfnI was the only functional LTA-synthase (**Fig 4A**). From this analysis, short LTA (<10 kDa) were more abundantly produced in NB-glucose. This seemed to be related to LtaS activity, as we observed an increase in short LTA in *ΔyfnI* and *ΔyfnI ΔyqgS* mutants compared to 168CA (**Fig 4A** left panel, 3^rd^ & 5^th^ lanes). Collectively, our results show that the regulation of LTA synthesis is more complex than thought (18) as it influenced by the metabolic/nutritional status of the cells.

**Figure 4.**
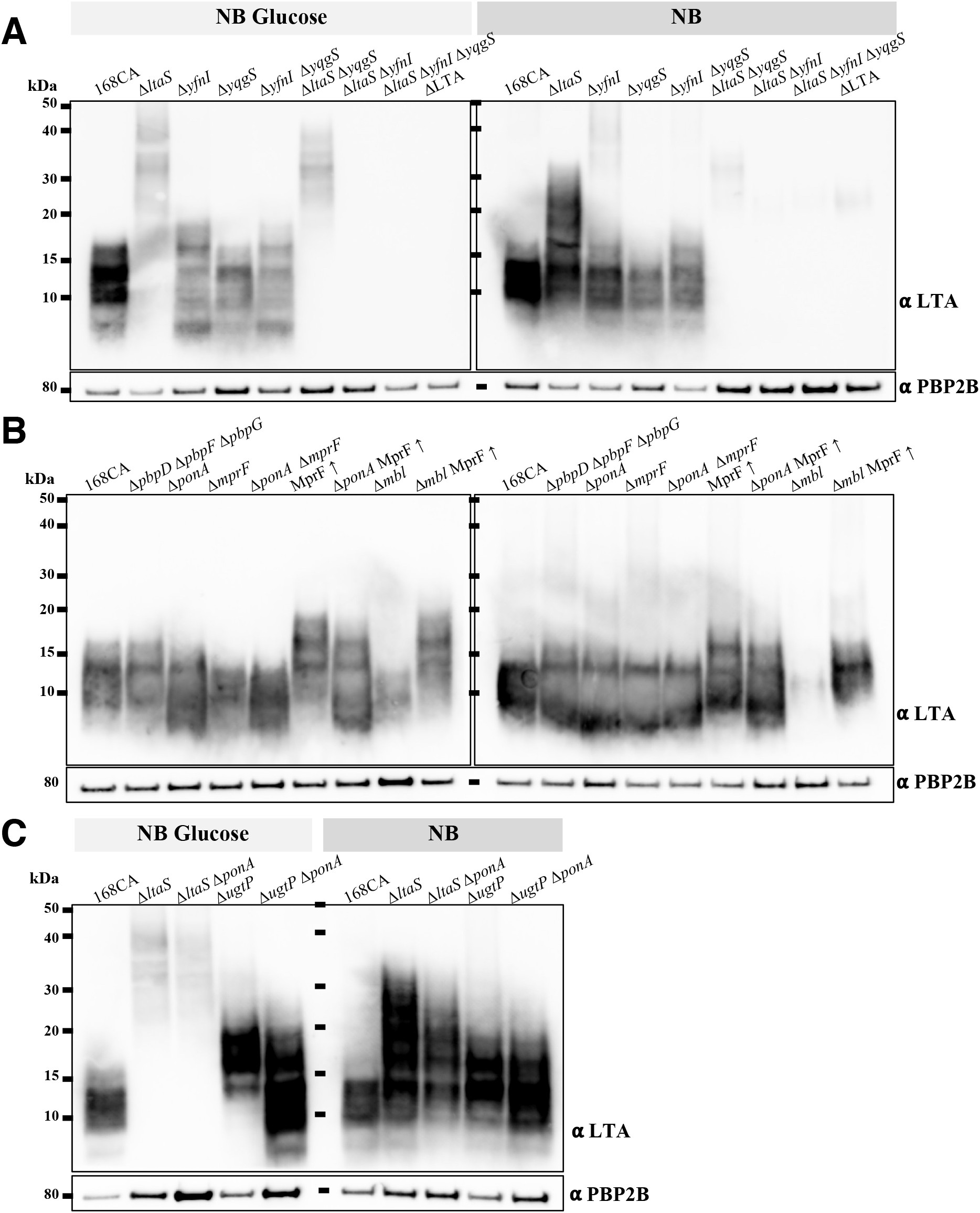
Analysis of lipoteichoic acid production in *B. subtilis* strains. For each set of cultures (NB ± Glucose 0.2%), samples were separated on the same Bis-Tris gradient gel. Membrane was split at ∼ 60 kDa position. The lower part of the membrane was probed using a monoclonal antibody for Gram-positive LTA and an HRP-linked anti-mouse antibody. The other membrane part was probed using polyclonal anti-PBP2B and HRP-linked anti-rabbit antibodies, to provide a sample loading control. **(A)** Samples were prepared from strains *B. subtilis* wild type (168CA), 4285CA (*ΔltaS*), 4289CA (*ΔyfnI*), 4292CA (*ΔyqgS*), AG595 (*ΔyfnI ΔyqgS*), AG593 (*ΔltaS ΔyqgS*), AG594 (*ΔltaS ΔyfnI*), AG600 (*ΔltaS ΔyfnI ΔyqgS*) and 4620 (*ΔltaS ΔyfnI ΔyqgS ΔyvgJ*, noted here ΔLTA). **(B)** Strains analysed: *B. subtilis* 168CA, AG417 (*ΔpbpD ΔpbpF ΔpbpG*), RE101 (*ΔponA*), AG181 (*ΔmprF*), AG193 (*ΔponA ΔmprF*), AG304 (MprF 1’), AG311 (*ΔponA* MprF 1’), 4261CA *(Δmbl),* AG322 *(Δmbl* MprF 1’). A densitometry graph of the left blot is presented in **Fig S5D**. **(C)** Samples were extracted from *B. subtilis* 168CA, 4285CA (*ΔltaS*), AG383 (*ΔltaS ΔponA*), PG253 (*ΔugtP*) and AG444 (*ΔugtP ΔponA*) strains. Data information: Representative of one of three independent experiments shown in **Fig S4-S6.** At each section A, B, C, samples were derived from the same experiment.

As the *mprF* mutation suppressed the *ΔponA* phenotype (**Fig 2A, Fig S1B**) we reasoned that any alteration of LTA synthesis would be detectable in a strain where only the major aPBP was deleted. As the multiple aPBP deletion strain was very lytic and did not grow under certain conditions (**Fig S1**), further experiments were done in the *ΔponA* background. Using this background, strains could grow in NB glucose 0.2% and this permitted the preparation of samples for the Western blot analysis. Analysis of *ΔponA* cells grown in the presence of glucose revealed that there was an increase in short LTA that was not evident in the absence of glucose (**Fig 4B** left panel, 3^rd^ lane, **Fig S5D**). In comparison, the LTA signal of *ΔponA ΔmprF* decreased compared to *ΔponA* (**Fig 4B** left panel, 5^th^ lane) and the *ΔmprF* mutant in NB-glucose (**Fig 4B** left panel, 4^th^ lane) decreased compared to that of 168CA (**Fig S5D**). Surprisingly, we found that *mprF* overexpression (MprF t) had a significant effect, resulting in an increased LTA polymer length (6^th^ lane in **Fig 4B)**. This effect was independent of YfnI, as a similar LTA profile was observed in a strain *ΔyfnI* MprF t while the profile of a *ΔltaS* MprF t was similar to that of *ΔltaS* (data not shown, strains AG353 & AG349 in **Table 2**). The latter was expected as LtaS activity seems dominant over YfnI (18). The overexpression of *mprF* in *ΔponA* cells resulted in a broader size range of LTA than that seen for the wild type as we observed an increase in LTA signals below 10 kDa but also above 15 kDa (7^th^ lane in **Fig 4B, Fig S5D**). This to some extent resembled that observed in single mutants *ΔponA* and MprF t, respectively. Thus, the cumulative effect of the mutations had led to an increase in LTA length range and could explain the lethality of the *ΔponA* MprF t double mutant on NA-glucose-IPTG (**Fig 2B**).

Western blot analysis of the LTA in the *Δmbl* background showed that it was weak, and representative of short LTA (second to last lane **Fig 4B, Fig S5D**). Overexpression of *mprF* rescued *Δmbl* growth (**Fig 2B**) and this overexpression resulted in an increased in LTA length and abundance in the *mbl* background (last lane **Fig 4B**). This result was also consistent with what was observed for *Δmbl ΔltaS* with the detection of longer LTA polymers (data not shown) similar to that of *ΔltaS* (**Fig 4A**). Thus an increase in LTA length seems to characterize the effect of these two *Δmbl* suppressor mutations.

Analysis of the suppressor *ΔltaS ΔponA* revealed the dominance of *ltaS* deletion, in that the LTA exhibited a retarded migration (3^rd^ lanes in **Fig 4C, Fig S6**), similar to that observed in *ΔltaS* (**Fig 4C**). In absence of UgtP which produces a lipid anchor Glc_2_DAG (**Fig 1**), we observed an increase in LTA length in both media conditions, but the shift was more distinct in glucose (>12 kDa, left panel 4^th^ lane in **Fig 4C**). The latter is comparable to the LTA signal detected in the *S. aureus ΔugtP* mutant grown in TSB, a glucose rich medium (32). Analysis of *ΔponA ΔugtP,* which was conditionally lethal on NA-glucose (**Fig S2C**), revealed an increase of the LTA-signals and length (7-25 kDa, left panel 5^th^ lane, **Fig 4C**), a pattern reminiscent of that determined for *ΔponA* MprFt strain (left panel 7^th^ lane in **Fig 4B)**, which was also lethal on NA-glucose (**Fig 2B**).

To conclude we show the first evidence that MprF activity has a role in altering the length of LTA, through a LtaS-dependent mechanism and has an impact on cell wall metabolism. Our results also highlight that the conditions leading to the lethality of *ponA* are associated with an increase in the abundance and length of LTA.

### Overexpression of major autolysin LytE is lethal in absence of PBP1 or LtaS

Our results inferred that the conditional lethality of aPBPs mutant might be linked to an interplay between LTA length and autolysin activity. Autolysins or cell wall hydrolases help shape and recycle the peptidoglycan during cell elongation and division. *B. subtilis* expresses a large number of them but only the loss of both PG D,L-endopeptidases CwlO and LytE is lethal (13, 14, 44–46) (**Fig 1A**). We focused on LytE because it is considered to be one of the key autolysins, it is secreted and thought to be anchored to the lateral cell wall and the septum in a way that is influenced by the teichoic acids (43). LytE abundance also increased in *ΔltaS* (47) and both LtaS and LytE affect colony development (48) and cell diameter (16, 43).

We reasoned that if an imbalance of LytE activity existed in the aPBPs mutants (*Δ4* and *ΔponA*) grown in glucose-rich medium, then a deliberate increase in LytE expression should exacerbate the strain phenotype, leading to cell death. Interestingly, *lytE* overexpression (by IPTG induction of a *P_hyspank_-lytE* allele) was found to be lethal in both *Δ4* and *ΔponA* backgrounds when grown on NA-glucose and it significantly delayed the growth of *Δ4* and *ΔponA* on NA but not that of *Δ3 ponA+* (**Fig 5A, Fig S7A**). This result confirmed that an imbalance in autolytic activity was detrimental to the cell that had lost PBP1. Importantly, the overexpression of *lytE* in the *Δ4 ΔltaS, ΔponA ΔltaS* and *ΔltaS* strains was lethal (**Fig 5A**) while the growth of *ΔponA ΔmprF* was only slightly delayed. However, the overexpression of *lytE* in *Δ4 ΔmprF* severely perturbed its growth on glucose but the strain remained viable. In contrast, the control strain with only the *ΔmprF* mutation was unaffected by *lytE* overexpression.

**Figure 5.**
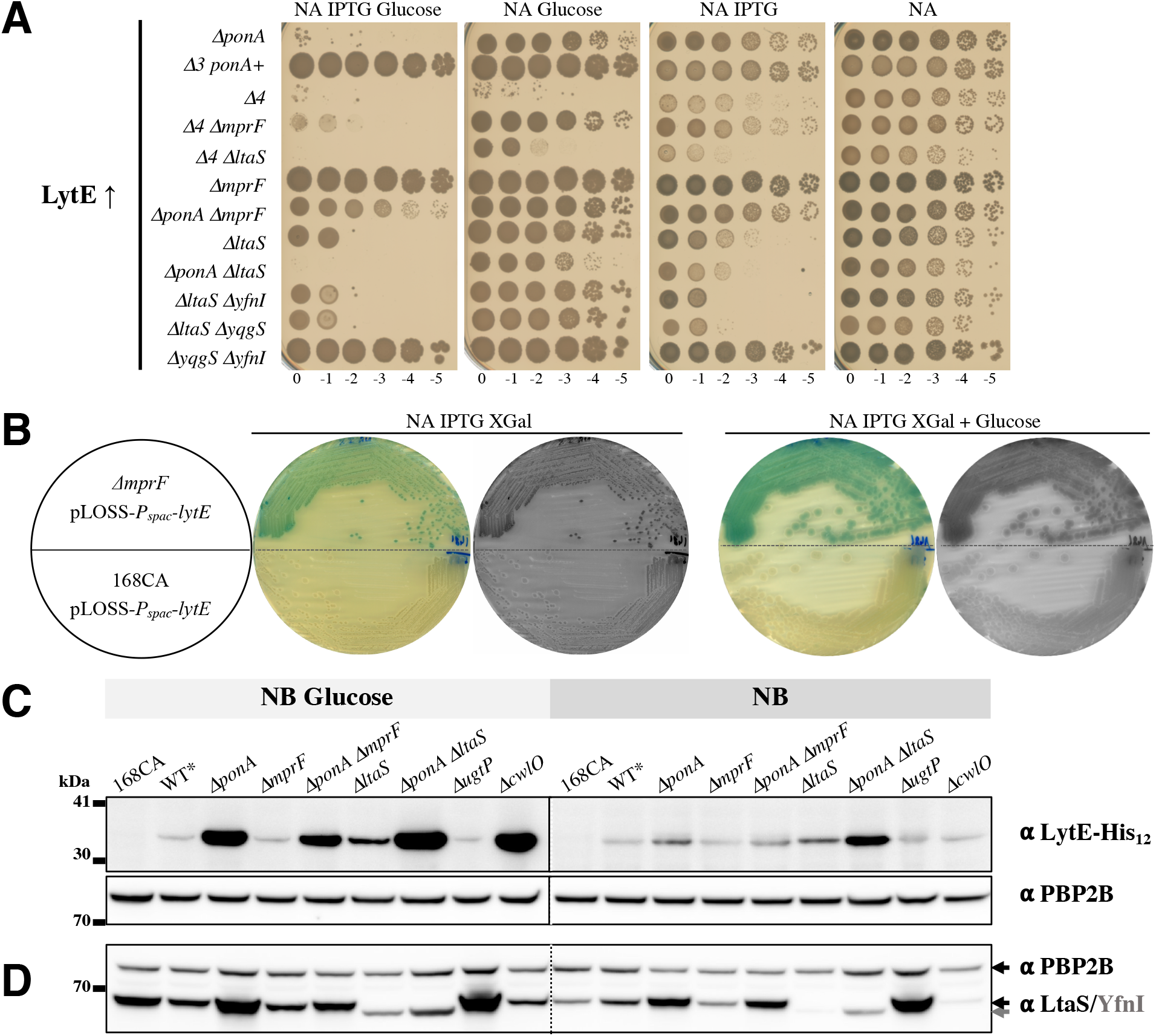
LytE overexpression is conditionally lethal in absence of PBP1 or LtaS. **(A)** *lytE* overexpression was analysed in strains which integrated at the *amyE* locus an (0.1 mM) IPTG-inducible *P_hyspank_-lytE,* represented here LytE t. The strains tested had the following relevant features: AG484 (*ΔponA* LytE t), AG1460 (*Δ3 ponA^+^* LytE t), AG501 (*Δ4* LytE t), AG502 (*Δ4 ΔmprF* LytE t), AG1465 (*Δ4 ΔltaS* LytE t), AG1478 (*ΔmprF* LytE t), AG1479 (*ΔponA ΔmprF* LytE t), AG1462 (*ΔltaS* LytE t), AG1480 (*ΔponA ΔltaS* LytE t), AG1497 (*ΔltaS ΔyfnI* LytE t), AG1498 (*ΔltaS ΔyqgS* LytE t) and AG1499 (*ΔyqgS ΔyfnI* LytE t). The following strains that did not display any obvious phenotype are not represented here: AG475 (168CA LytE t), AG1461 (*ΔlytE* LytE t), AG1463 (*ΔyfnI* LytE t), AG1492 (*ΔyqgS* LytE t). Plates (here glucose 0.5%) were incubated at 37°C and scanned after 24 h (this figure) and 48 h (**Fig S7A**). **(B)** Strains AG1481 (*ΔmprF* pLOSS-*P_spac_-lytE*) and AG551 (168CA pLOSS-*P_spac_-lytE*) were streaked on NA and NA glucose (0.5%), supplemented with IPTG (1 mM) and X-Gal from a fresh colony on NA spectinomycin. Plates were incubated at 37°C and scanned after 24 h. Blue colonies indicate the stabilisation of plasmid pLOSS-*P_spac_-lytE* in *ΔmprF*. Grey scale images are here to help visualize white colonies. **_(C)_** Detection of LytE-His_12_ and PBP2B by Western blots. For simplification on this figure only the relevant background feature is displayed for the following strains (expressing LytE-His_12_ under the control of its native promoter): wild-type-like AG565 (WT*), AG587 (*ΔponA*), AG1486 (*ΔmprF*), AG1487 (*ΔponA ΔmprF*), AG1489 (*ΔltaS*), AG1488 (*ΔponA ΔltaS*), AG1535 (*ΔugtP*) and AG1541 (*ΔcwlO*). In addition, *B. subtilis* 168CA was grown in parallel (without antibiotic) and used here as a negative control for the LytE-blot. The membrane was split and incubated either with a monoclonal Penta-His and an HRP-linked anti-mouse antibodies or with a polyclonal anti-PBP2B and HRP-linked anti-rabbit antibodies. Note: a ‘ladder’ sample loaded after the NB-Glucose (0.2%) sample set is not displayed and the image editing is symbolised by the vertical black line (see original on **Fig S7B,** middle panel). **(D)** Detection of LTA-synthases LtaS/YfnI and PBP2B by Western blots. The samples used in experiment **C** were loaded on a new SDS-gel and the Western blot was carry out with polyclonal anti-LtaS (able to detect YfnI, unpublished Errington’s lab), anti-PBP2B antibodies, and HRP-linked anti-rabbit antibodies. Data information: One representative set of three independent Western Blot experiments is shown here and in **Fig S7B, S8A**. Additional supporting information is presented in **Fig S8, S9**. At each section samples were derived from the same experiment.

This finding suggested that *ΔponA ΔltaS* and *ΔponA ΔmprF* mutants, which have distinct LTA length, were differently sensitive to LytE activity. Therefore, we examined the double LTA-synthase mutant backgrounds, here we found that only the strain expressing *ltaS* was viable when *lytE* was overexpressed (**Fig 5A,** last strain). In these genetic backgrounds, the absence of *mprF* or *ugtP* had no effect (data not shown). Interestingly, *ΔltaS* LytE t was viable on NA (**Fig 5A,** 8^th^ strain) but not on NA-glucose while we previously identified that *ΔltaS* produces distinct LTA length on each of these media (**Fig 4A**). Thus, it is clear that the activity of LtaS, is required for the cell to balance the effect of overexpression of *lytE*, presumably by producing a specific LTA length.

To comprehend the apparent tolerance of LtaS^+^ and *ΔmprF* cells to LytE, we constructed an unstable replicon pLOSS-*P_spac_*-*lytE* plasmid which expresses *lytE* at a level lower than *P_hyspank_-lytE* (**Table 1**). This plasmid in the *ΔmprF* strain was retained in all media conditions tested (AG1481 strain), including elevated magnesium. This was indicated by the fact that the colonies remained blue on IPTG XGal supplemented media (**Fig 5B**) whereas other strains tested showed instability of the plasmid (e.g. 168CA, *ΔponA, ΔponA ΔmprF* and LTA-synthases mutants, **Table 2**) an indication that there was no selective advantage for the cells to maintain the plasmid. Thus this plasmid stability assay highlights that *ΔmprF* cells tolerate higher expression of *lytE* compared to the other background strains tested. At this stage, it is unclear if the *ΔmprF* strain’s tolerance for *lytE* overexpression is due to the loss of aaPGols or the production of shorter LTA or both.

### LTA influences LytE activity

The conditional lethality observed when overexpressing *lytE* prompted us to investigate the cellular levels of this autolytic enzyme in our strains. First, we engineered strains that could express a recombinant protein LytE-LEMGRSH_12_ (using pMUTin-*‘lytE-his,* **Table 1**) and then analysed total lysates by Western blot using anti-histidine (for LytE) (**Fig 5C, Fig S7B**) and an anti-LtaS antibody raised against LtaS (74 kDa) that cross reacts with YfnI (73 kDa) (**Fig 5D, Fig S8A**). This showed that LytE and LtaS/YfnI accumulate in *ΔponA* cells, particularly when grown in presence of glucose (**Fig 5C-D**). Using a set of strains that had either pMUTin-*‘ltaS-his* or pMUTin-*‘yfnI-his* integrated into the genome, we could confirm that both LtaS and YfnI are more abundant in *ponA* cells (**Fig S8B, Tables 1-3**).

In *ΔponA ΔmprF*, LytE and LtaS/YfnI protein levels were lower compared to *ΔponA* (**Fig 5C-D**), but no obvious change was observed in the wild-type background (WT*) and *ΔmprF* strains. *ΔltaS* and *ΔponA ΔltaS* exhibited an increase in both LytE and YfnI when grown in NB-glucose, an effect more pronounced in the *ΔponA ΔltaS* mutant (**Fig 5C-D**, left panel). In addition, comparative analysis of the set of LTA-synthase mutants (**Fig S8C**) revealed that LtaS/YfnI increased in presence of glucose and in the absence of YfnI, LtaS increased significantly. Also, and potentially more importantly, in the absence of LtaS, LytE abundance increased. Analyses of the strains that were lytic under the same growth conditions showed that there was an excess of LtaS/YfnI in *ΔponA ΔugtP* and *ΔponA* MprF t cells cultured in either NB or NB-glucose (**Fig S9B-C,** e.g. 6-7^th^ lanes). It was also evident that LytE was more abundant than the wild type. These observations seem to support the idea that the altered abundance of these proteins, identified in *ΔponA* cells, was associated with the sick phenotype.

Taken together, our results show a correlation between LTA length and LytE abundance in the cell which suggests that LTA acts to moderate cell wall metabolism. In *ponA* mutants, this function is unbalanced such that cell wall degradation is predominant. This is summarised in the discussion in **Fig 7**.

**Figure 6.**
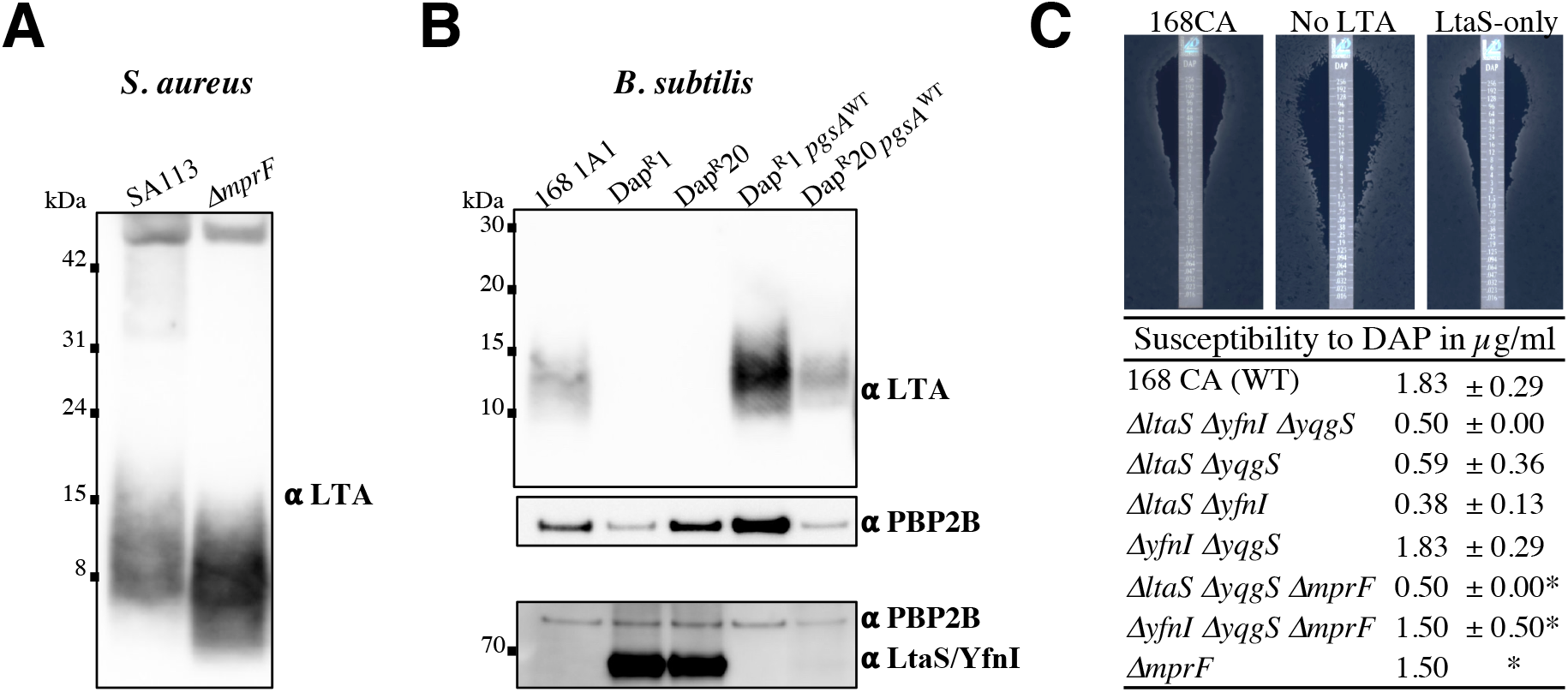
Daptomycin resistance and LTA synthesis in *B. subtilis* and *S. aureus*. **(A)** Samples of methicillin-sensitive *Staphylococcus aureus* strains SA113 and SA113 *ΔmprF* (21) were probed for LTA by Western Blot. **(B)** Researchers identified that a modified allele of *pgsA*(*A64V*) was the major determinant of daptomycin resistance of *B. subtilis* Dap^R^1 and Dap^R^20 strains selected by serial daptomycin antibiotic passages (54). The strains 168 1A1 (wild type, WT), Dap^R^1, Dap^R^20, HB15516 (Dap^R^1 *pgsA*^WT^) and HB15507 (Dap^R^20 *pgsA*^WT^) were grown in NB overnight at 30°C and LTA was detected (top panel) along with PBP2B (middle panel) by Western Blots. Analysis of the samples grown exponentially at 37°C using theses overnight cultures is presented in **Fig S10F**. The bottom panel shows the cellular abundance of the LTA-synthase detected with an anti-LtaS (that also cross reacts with YfnI, unpublished Errington’s lab). Here the samples were normalized based on the previous PBP2B blot (middle panel) (blotted as in **Fig 5D**). Data information: One representative set of three experiments is shown here and in **Fig S10F**. **(C)** Absence of LTA-synthase LtaS increase *B. subtilis* susceptibility to daptomycin. Dap strip assays (Liofilchem) on NA-CaCl_2_ plates were carried out on *B. subtilis* wild type (168CA), AG600 (*ΔltaS ΔyfnI ΔyqgS,* noted no LTA), AG593 (*ΔltaS ΔyqgS*), AG594 (*ΔltaS ΔyfnI*), AG595 (*ΔyfnI ΔyqgS,* noted LtaS-only), AG1673 (*ΔltaS ΔyqgS ΔmprF*), AG1675 (*ΔyfnI ΔyqgS ΔmprF*) and AG1663 (*ΔmprF*) strains. Plates were incubated at 37°C and scanned after 24h (illustrations on top panel). * The table represents the average of at least two independent experiments except for *ΔmprF*. Data information: Additional supporting information is presented in **Fig S10**

**Figure 7.**
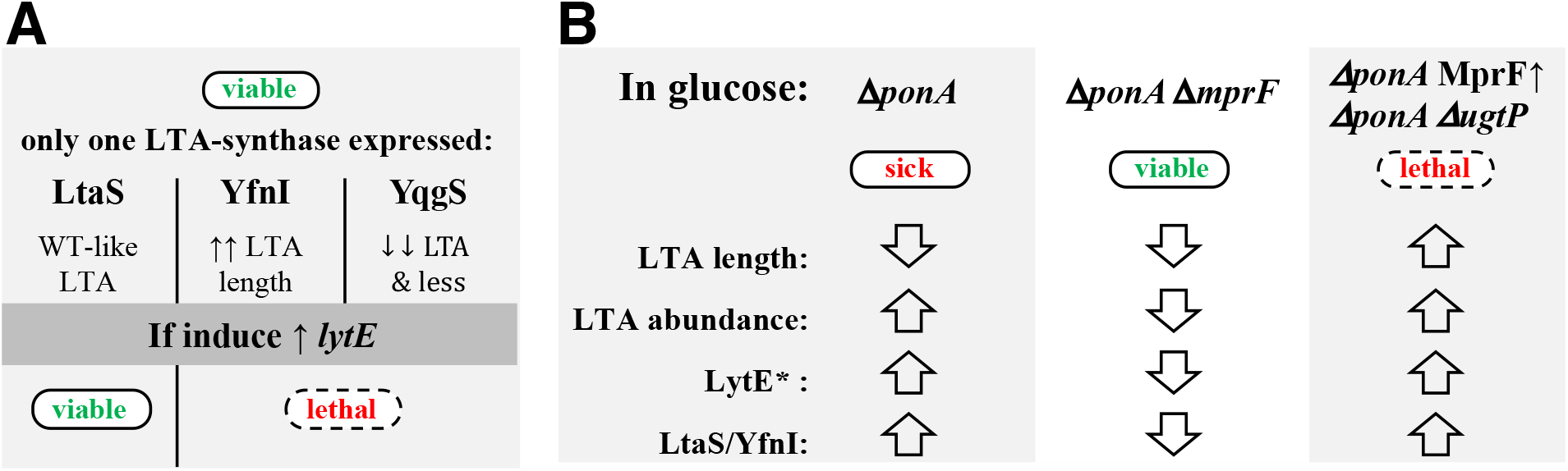
A summary of the alteration in LTA and LytE levels associated with the conditional lethality of *ΔltaS* and *ΔponA* mutants. **(A)** The overexpression of *lytE* (inducible) in the double LTA-synthase mutants is lethal except when the major LTA synthase LtaS is functional. The presence of a distinct LTA length is presented by the mutant strains (**Fig 4A, 5A**). **(B)** In glucose, the viability and lethality of mutants in *ΔponA* background were associated to distinct LTA, LytE and LtaS/YfnI levels (**Fig 2A-B, 3A, 4B-C, 5A, C-D**). * The overexpression of *lytE* (inducible) is lethal to *ΔponA* but not to *ΔponA ΔmprF* mutant (**Fig 5A**). Strains with the ability to grow without significant cell lysis are referred as viable (green text in the cell). Lethal refers to an absence of growth of the mutant on NA or in the specified condition (dashed rod cell, with red text). In these cases, analysis of the strains was done by shifting cultures to non-permissive conditions (e.g. low magnesium, addition of glucose or inducer). Arrows indicate an increase (τ) or decrease (−ϑ) of the specified factor. Here it is assumed that the strength of LTA signal observed by Western blot is proportional to abundance.

### LTA production is altered in *S. aureus ΔmprF* and is an important factor in *B. subtilis* sensitivity to daptomycin

MprF is conserved in the *Firmicutes* and has been shown to contribute to *S. aureus* pathogenicity (21, 22). Importantly, studies of clinical isolates have found that methicillin-resistant *S. aureus* strains (MRSA) that have acquired resistance to daptomycin (Dap^R^), a lipopeptide antibiotic that acts to interfere with membrane containing phosphatidylglycerol (26, 49–51), frequently carry nucleotide polymorphisms (SNPs) in *mprF* (52, 53). Although some *mprF* SNPs were associated with a MprF gain of activity (increasing LysPGol and reducing PGol), others had no clear correlation between the phospholipids composition and the relative surface charge of the cell (26). In this context, we asked whether the role of MprF in LTA production is conserved in *S. aureus* as it seemed reasonable that this might contribute to Dap resistance. Analysis of LTA production in methicillin sensitive *S. aureus* strains showed that the LTA from the *mprF* mutant migrated faster than that of the wild type SA113 (**Fig 6A, Fig S10A**), thus LTA production is altered in *S. aureus ΔmprF*. Interestingly, when exposing exponentially growing cells of *S. aureus* to daptomycin, our analysis showed that the LTA length and production had increased in both strains (**Fig. S10B-C**, samples treated with 0.25 and 0.5 μg/ml DAP for *ΔmprF* and the wild type, respectively). It is also noteworthy that even when exposed to DAP, the LTA produced by the wild type remained longer than that of the *ΔmprF* mutant, which perhaps indicates that the LTA length plays a role in DAP sensitivity. This effect was observed at DAP concentrations that did not significantly impacted on the wild type growth (**Fig. S10E**) whereas the growth of the *mprF* mutant was significantly impaired (**Fig. S10D-E**), as previously reported (52, 53). Our results seem to confirm that MprF’s role in modulating LTA production is conserved in both *Bacillus* and *Staphylococcus* species. Collectively, these results indicate that a change in LTA nature and/or production, rather than an alteration in membrane properties, might be associated with Dap resistance in *S. aureus*.

As we could not carry analysis on Dap^R^ MRSA (laboratory regulations), we, therefore, turned our interest to the strongest known Dap^R^ allele in *B. subtilis, pgsA A64V* (54). This mutation of the essential gene *pgsA,* encoding the PGol-synthase (**Fig 1B**), has been shown to reduce cellular PGol content and was suggested to modify LTA production (54). Thus, we analysed the LTA of *B. subtilis* wild type (WT), isogenic strains carrying *pgsA^A64V^* (strains Dap^R^ 1 & Dap^R^ 20) and *pgsA^A64V^* complemented with *pgsA*^WT^ grown overnight in NB **(Fig 6B, Fig S10F)**. Our assays revealed that no LTA signal was detected below 30 kDa in Dap^R^ strains whereas LTA was detected in the strains complemented for *pgsA* (Dap^R^ *pgsA*^WT^) that were similar to that of the wild type strain. Thus, PgsA A64V accounts for the severe reduction in LTA production. Finally, we verified that the two major LTA-synthases were still produced in Dap^R^ strains by Western blot (**Fig 6B, Fig S10F**). A strong LtaS/YfnI signal was detected in Dap^R^ strains which proved that one or both LTA-synthases were still expressed, but up-regulated.

The above findings in *S. aureus* and *B. subtilis* suggested that alteration in LTA could be associated with DAP sensitivity. As *S. aureus ltaS* is essential, we decided to analyse the Dap-sensitivity of *B. subtilis* expressing only one LTA-synthase. Our assays on NA (**Fig 6C**) and NB (**Fig. S10G**) showed that strains lacking the major LTA-synthase LtaS were sensitive to DAP and this sensitivity was higher than that observed for *ΔmprF* or *Δdlt* mutants (**Fig S10G**). Thus LtaS activity is required for DAP tolerance in *B. subtilis*, more so than MprF.

## Discussion

In absence of the class A PBPs (*11.4*), *B. subtilis* is viable due to the presence of RodA, the essential PG glycosyltransferase (controlled at least in part by the cell envelope stress regulator σ^M^ (11, 12)) and yet lethal on glucose rich medium (**Fig 2A** & **S1B**). By studying either *11.4* or *11.ponA* single mutant, we now understand that the presence of glucose leads to an increase in abundance of the LTA-synthases LtaS and YfnI. The increase in these enzymes results in altered LTA production that in turn causes the accumulation of major autolysin LytE (**Fig 4-5**), the end result being cell lysis through the inability to regulate the activity of autolysins (**Fig 7**). This lytic phenotype could be suppressed by the absence of MprF (**Fig 2A, S1B**), an enzyme that we think helps in the regulation of LTA production (**Fig 8**). In *11.ponA,* loss of MprF resulted in reduced LTA production and decreased level of LytE (**Fig 4B, 5C, S5D**). This was consistent with our finding that an increase in length and abundance of LTA (overexpression *mprF*, *11.ugtP*) and/or an increase in LytE correlated with glucose-mediated lethality in *11.ponA* (**Fig 2B, 3A**, **4B-C, 5A, S5D, Fig S9B-C**). Loss of major LTA-synthase LtaS was not as beneficial for the growth of *11.4* as the loss of MprF (**Fig 3B, S1B**). In *11.ponA,* loss of LtaS lead to the production of very long LTA polymers similar to the single *ltaS* mutant and a persisting high level of LytE (**Fig 4C, 5C**). This might explain why the growth of *11.ponA 11.ltaS* was not significantly enhanced compared to *11.ponA* (**Fig S1B**). In support of this, the analysis of strains only defective in LTA synthesis (last three rows in **Fig 5A**) indicated that LtaS presence is required to prevent cell lysis when *lytE* was artificial overexpressed (**Fig 5A, 7**), presumably because a distinct LTA length (7-17kDa) and/or abundance similar to the wild type LTA (**Fig 4A**) is required (**Fig 7**).

**Figure 8.**
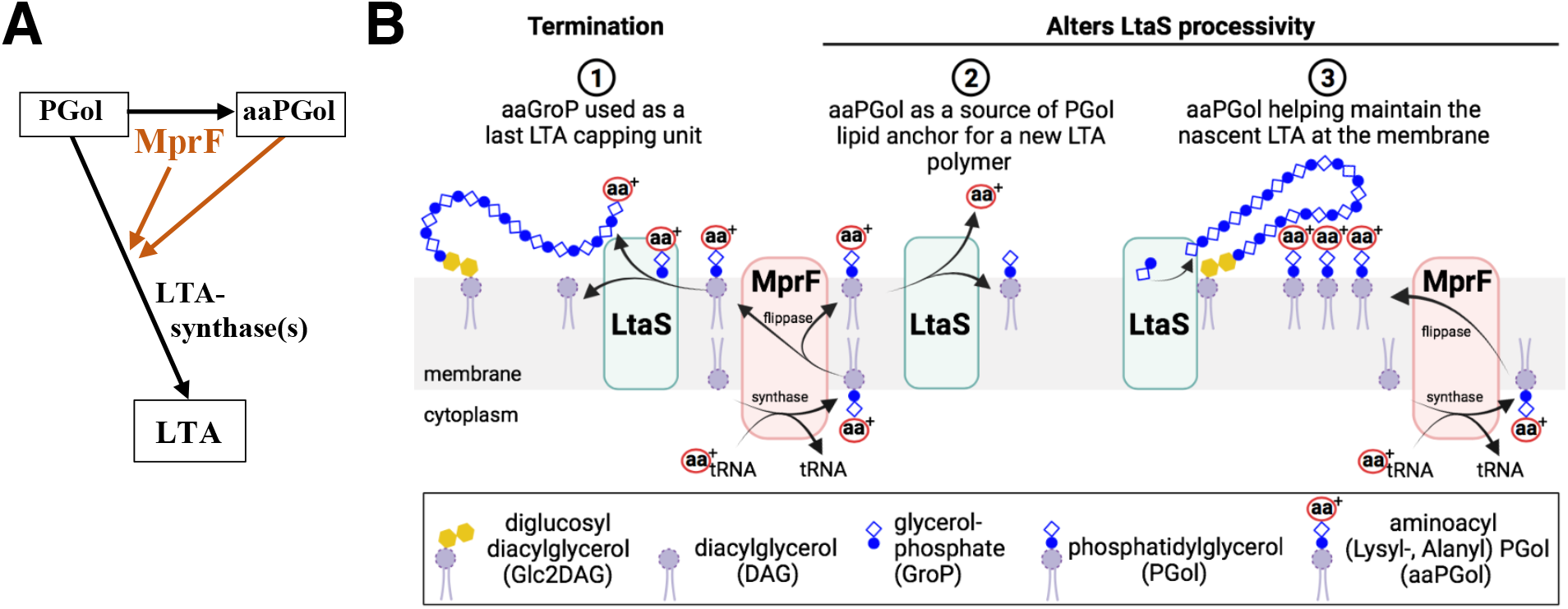
MprF positively regulates the synthesis of LTA in *B. subtilis* and *S. aureus*. (**A**) MprF or its substrates positively regulate LTA production in *B. subtilis* and in *S. aureus*. (**B**) Proposed role of MprF in LTA production via its substrate aminoacyl-phosphatidylglycerols (aaPGol) (not limitative). (**1**) LtaS could use the aminoacyl-glycerol-phosphate (aaGroP) of aaPGol as a last LTA capping unit. Potentially the addition of this positive charged capping unit (e.g. lysyl, alanylGroP) could contribute to the protection against CAMPs. (**2-3**) MprF could indirectly alters LtaS processivity: the release of the aminoacyl could permit LtaS to use the PGol as lipid anchor (**2**), the presence of aaPGol in the vicinity of LtaS could create an electrostatic environment that help maintain the growing LTA chain in a conformation that favors LtaS processing (**3**).

Previous studies have suggested that LTA could inhibit autolysins activity (1, 5) but the key factors involved have not been identified. Recent work in *S. aureus* showed a *11.ugtP* mutant producing long LTA is susceptible to PG hydrolases (55) and it has been suggested that major autolysin LytA is sequestered by LTA in exponential growth (56). Our results indicate that LytE in *B. subtilis* exhibits similar characteristics and that the length of LTA has a role in modulating LytE abundance or presumably its activity (**Fig 7**). However it is unclear how this mechanism may work in relation to the localisation of the LTA in the wall relative to active autolysins, particularly LytE, and this remains an important question to answer in future work.

In *B. subtilis* 168, expression of *lytE* is quite complex and coordinated by regulators such as α^I^ and WalR (46, 57). In contrast, transcriptional regulation of *mprF* is not well defined, but interestingly analysis of *B. subtilis* BSB1 showed that *mprF* and *pgsA* expression is positively correlated with that of *walKR,* the two-component response system that controls cell wall metabolism, and notably *lytE* (cluster A418 in Subtiwiki expression data browser; (58, 59)). However, expression of *walKR* and *sigI* is not significantly altered in LB-glucose (58) or in the absence of aPBPs (12) but theses regulators might play a post-transcriptional role in *11.4* where the α^M^ cell wall stress response is active (12, 57). This seems to be supported by the fact that some of the cell wall associated genes mentioned (*sigM*, *ponA*, *ltaS*, *yfnI*, *mprF*, *pgsA*) are differently expressed in cells of *B. subtilis* BSB1 in LB-glucose (58, 59). Thus *B. subtilis* might need to adapt its cell wall metabolism in response to glucose and this might account for the partial redundancy of function of *B. subtilis* LTA-synthases. Evidence also suggest that the presence of magnesium in culture medium leads *Bacillus* cells to adjust its cell wall metabolism. When we looked at the effect of magnesium in presence of glucose on aPBP mutants, as this is known to suppress the growth defect (10), we have observed that magnesium decreased the abundance of LytE significantly in *ΔponA* and *ΔponA ΔmprF* (**Fig S9**) but not as much in *ΔponA ΔltaS.* In addition, high levels of magnesium have an impact on LTA production (work in progress) and other processes (60, 61). Thus, the mechanism by which exogenous magnesium helps the growth of some cell wall mutants might partially require LtaS activity to tune LytE abundance in the cell wall (14). Altogether, our study brings a new perspective on the importance of LTA in the cell envelope stability and how the extracellular biosynthesis is regulated.

In this study, we have identified a new function for MprF, an enzyme known to synthetise aminoacyl-phosphatidylglycerol (aaPGol) phospholipids and translocate them across the membrane (**Fig 1**). We show that MprF activity impacts the biosynthesis of LTA (**Fig 8A**) and this in turn alters cell wall metabolism. The LTA profiles observed in *B. subtilis* and *S. aureus* lacking MprF (seen as shorter and/or decreased in abundance) and the overexpression of *B. subtilis* MprF (causing a smeared & retarded mobility of the LTA) (**Fig 4B, 6A**) suggest that MprF positively regulates LTA production (**Fig 8A**). Other studies have indicated that MprFs may have relaxed substrate specificities across species (22, 62) and even synthetise lysyl-glucosyl-DAG glycolipid in *Streptococcus agalactiae* (63). Recently, it has been shown that when *S. aureus* LtaS uses PGol as an alternative anchor to the glycolipid produced by UgtP, the concentration of these free lipid starter units regulates the length of the LTA polymerised *in vitro* (33). Thus it is possible that MprF activity contributes to LTA synthesis (**Fig 8B**). MprF might provide the last capping unit (Lys-, Ala-GroP) to LtaS to terminate the LTA being polymerised, this would also contribute to the protection against CAMPs (**Fig 8B**). Alternatively, aaPGols might be a source of GroP units (**Fig 8B**) in the formation of LTA in addition to the known PGol source altering processivity (33). Another possibility would be that aaPGols change the electrostatic environment and this impacts on LTA-synthase(s) activity by keeping the LTA polymers adjacent to the membrane (**Fig 8B**).

Gain of function in MprF has been identified in methicillin-resistant *S. aureus* (MRSA) clinical isolates that have acquired resistance to daptomycin (Dap), an antibiotic used as a last resort to treat MRSA infections (26, 53). Over the years, MprF-mediated resistance to Dap (Dap^R^) has been investigated but a consensus model has yet to link Dap^R^ to a particular domain of MprF and/or to a particular LysPGol content in one or both membrane leaflets (26, 64). In *B. subtilis,* our analysis shows the significant decrease of LTA produced in a strain with the *pgsA A64V* mutation (**Fig 6B**), a strain with a reduced PGol content that exhibits Dap resistance (54). Also, we found that LTA-synthase LtaS is required for DAP tolerance in *B. subtilis* (**Fig 6C**). The function of MprF in LTA synthesis and the new understanding of *S. aureus* LtaS processivity (33) could be the link with the alleles *cls* encoding a cardiolipin synthase, *pgsA* and *dlt* operon (50) identified in *S. aureus* MRSA that result in Dap resistance. Together, these findings support the idea that a modification in LTA polymerisation in addition to D-alanylation of teichoic acid (65, 66) could result in a cell envelope that is less permeable to Dap, and not necessarily linked to increased cell wall thickness (67). Further analysis of LTA synthesis and cell wall composition will help redefine MprF-mediated Dap^R^ mechanism and help appreciate the exact role of MprF in the net cell surface charge and the protection against cationic antimicrobial peptides. Recent research on compounds targeting LTA synthesis (68, 69) and on MprF-targeting monoclonal antibodies (70) suggest these approaches have relevance in combating clinically relevant bacteria.

## Material and Methods

### Plasmids, bacterial strains, and primers

Plasmids and strains used in this study are listed in **Tables 1** and **2.** Strains constructions are described below or in **Table 2** (last column). Primers used are listed in **Table 3.**

**TABLE 3.**
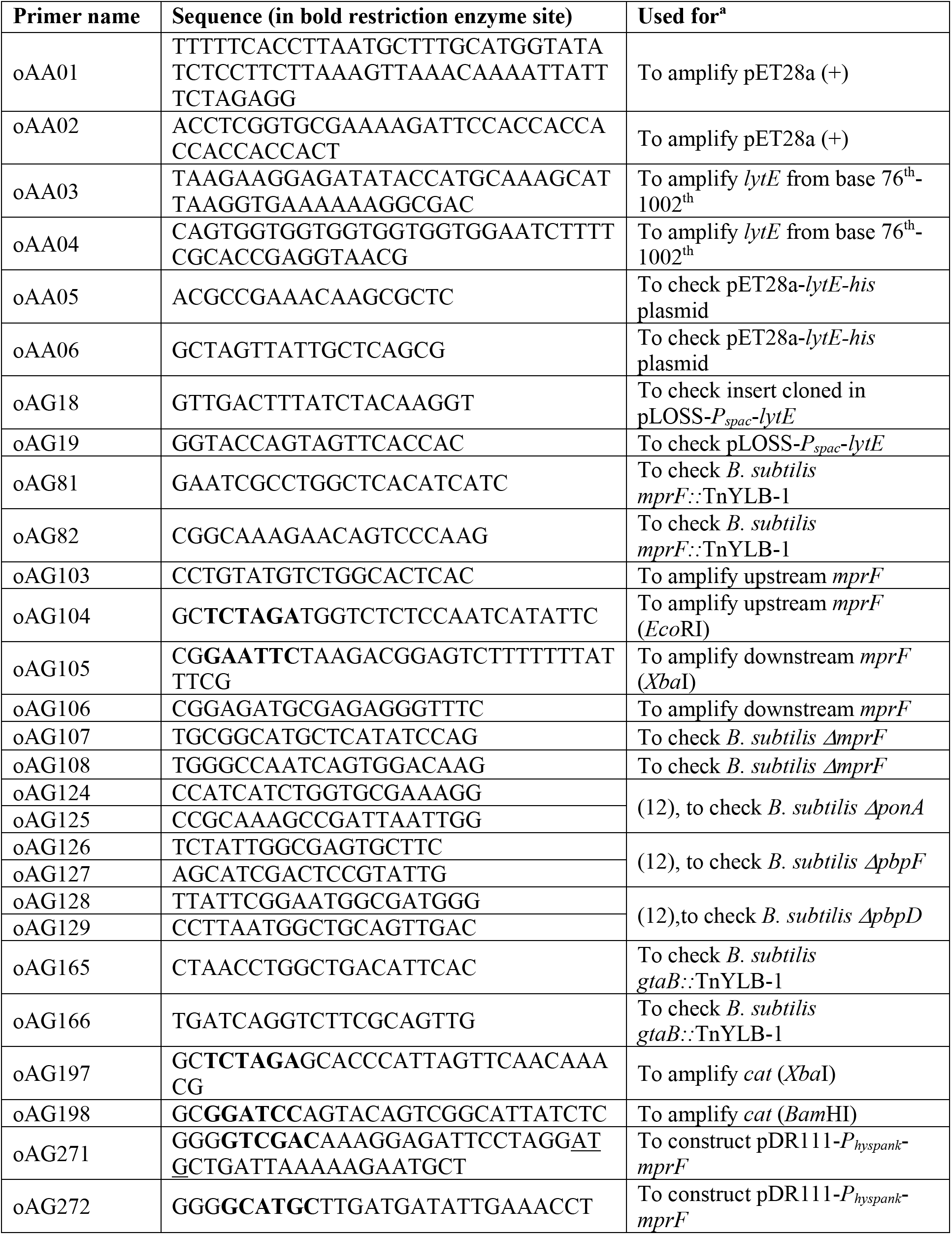
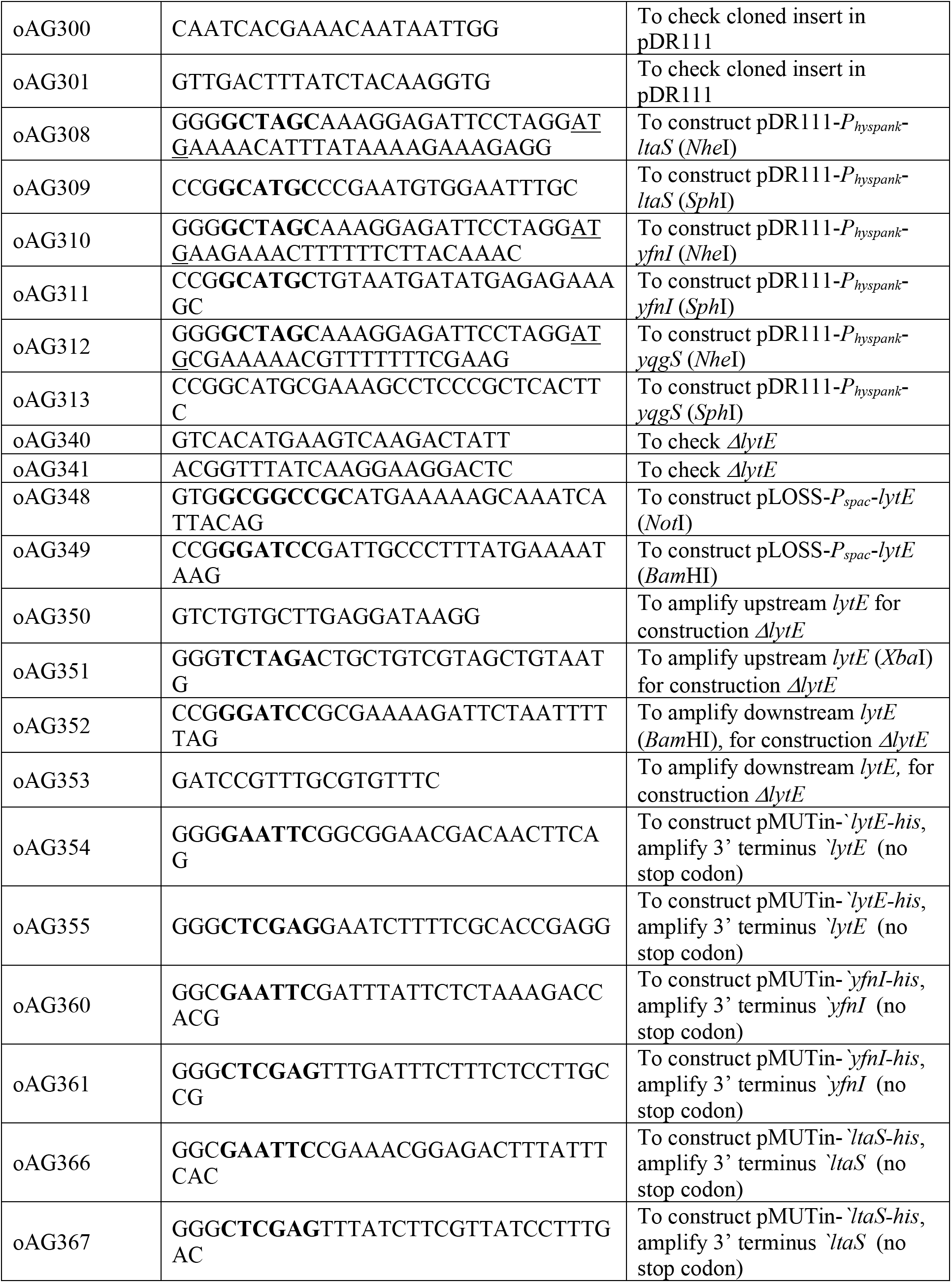
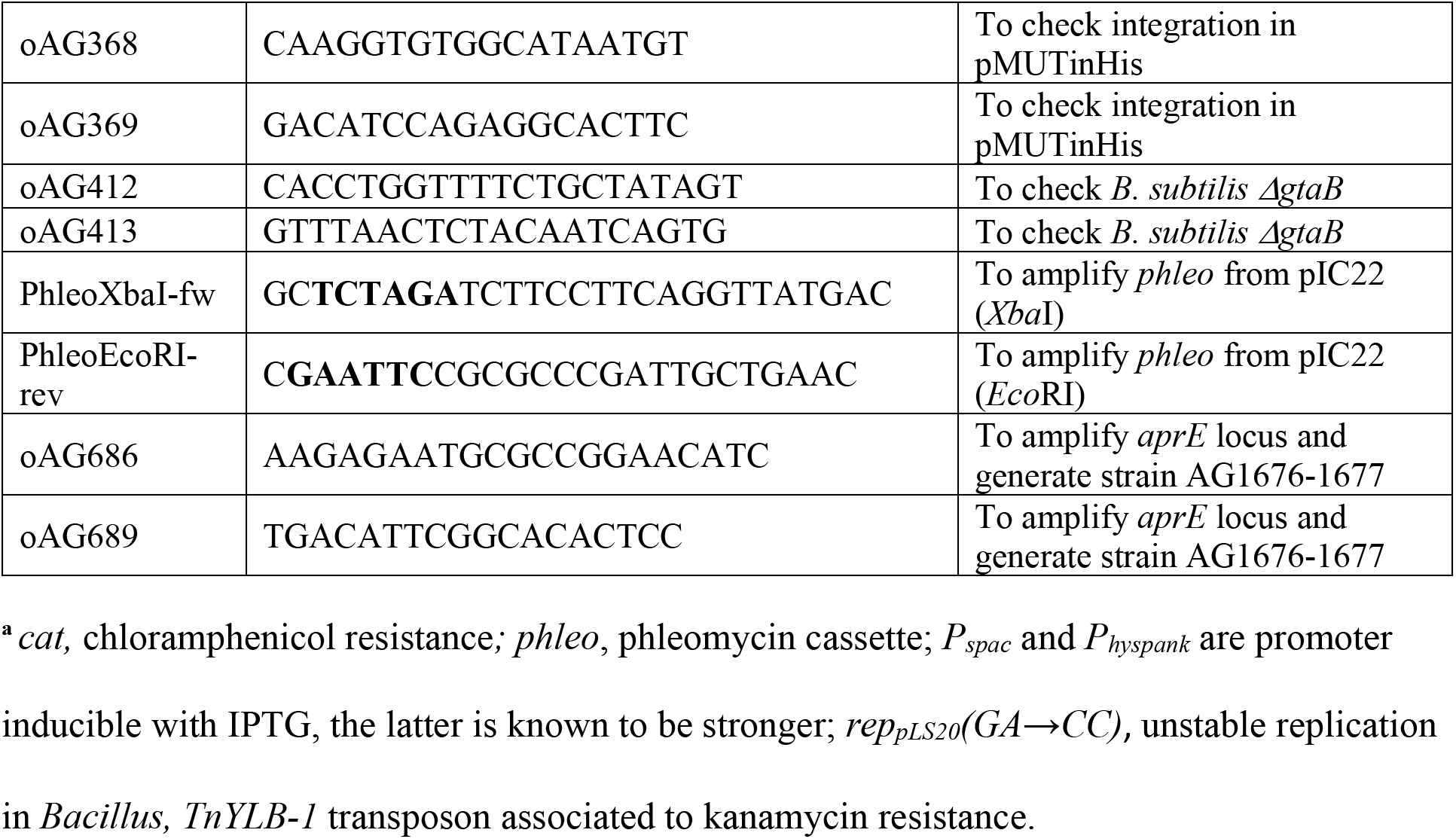
Primers used in this study

### Growth conditions and general methods

*B. subtilis* strains were grown routinely on nutrient agar (NA, Oxoid) or in nutrient broth (NB, Oxoid). For some assays, strains were grown in liquid Difco antibiotic medium 3 (PAB, Oxoid) and PAB agar (15 g/l agar, Oxoid). *B. subtilis* transformations were carried out as previously described using SMM defined minimal medium (71, 72). *B. subtilis* was usually incubated at 30°C (overnight), next day cells were diluted and grown at 37°C (unless specified differently). *Escherichia coli* and *Staphylococcus aureus* strains were incubated at 37°C on NA plate or in Luria-Bertani (LB). PCR, plasmid manipulations, and *E. coli* transformation were carried out using standard methods. Strains were selected on NA supplemented with ampicillin (for *E. coli,* 200 μg/ml), spectinomycin (60 μg/ml), kanamycin (5 μg/ml), phleomycin (1 μg/ml), erythromycin (0.5 μg/ml), chloramphenicol (5 μg/ml) and daptomycin (see later). Various supplements were also used: IPTG (isopropyl-β-D-thiogalactopyranoside, 0.1 or 1 mM), X-Gal (5-bromo-4-chloro-3-indolyl-β-D-galactopyranoside, 100 μg/ml), MgS0_4_ (between 5 mM to 25 mM as required), and glucose (final concentrations up to 1%). For NA plates supplemented with 20 mM MgSO_4_, the concentration of spectinomycin, kanamycin and erythromycin antibiotics was doubled to avoid growth of false positive strains. Strains with the pMUTin*-his* integration were always grown in presence of erythromycin.

### Plasmids constructions

To obtain pDR111*-P_hyspank_-mprF* plasmid, the *mprF* (BSU_08425) coding sequence was amplified using primers oAG271-272, which changed the start codon (from TTG to ATG). The PCR product and pDR111 plasmid (**Table 1**) were digested by *Sal*I and *Sph*I restriction enzymes, ligated together and transformed into *E. coli* DH5α. Digestion and PCR sequencing using primers oAG300-301 verified the presence of the insert.

To construct pDR111-*P_hyspank_*-*ltaS*, pDR111-*P_hyspank_*-*yfnI* and pDR111-*P_hyspank_*-*yqgS,* each LTA-synthase gene was amplified by PCR with primers pairs oAG308-309 (2064 bp), oAG310-311 (2078 bp) and oAG312-313 (1990 bp), respectively. PCR products were digested by *Nhe*I and *Sph*I, each ligated with the plasmid pDR111 digested with the same restriction enzymes. The next steps were carried out as described for pDR111*-P_hyspank_-mprF*.

For pLOSS-*P_spac_-lytE, lytE* was amplified by PCR with oAG348-349 (1143 bp) primers, and the PCR product inserted in pLOSS* (**Table 1**) as a *Not*I and *Bam*HI fragment. After transformation, the resulting plasmid was checked by PCRs with primers oAG18-19 and finally checked by sequencing.

To construct the plasmids pMUTin-*‘lytE-his*, pMUTin-*‘ltaS-his* and pMUTin-*‘yfnI-his* the 3’-terminus part of gene of interest gene (except for the stop codon) was amplified with primer pairs oAG354-355, oAG366-367 and oAG360-361, respectively to obtain a PCR products ranging from 288 to 309 bp. The PCR products and the plasmid pMUTinHis (**Table 1**) (73) were digested by *Eco*RI and *Xho*I, ligated together and transformed into *E. coli* DH5α. Each insertion ‘*lytE, ‘ltaS* and *‘yfnI* was verified by PCR and plasmid sequencing using primers oAG368-369.

To construct pAM-21, plasmid pET28a (+) (Novagen) and *lytE* (from the 76^th^ bp, removing the coding sequence for the signal peptide) were amplified by PCRs using primer pairs oAA01-02 and oAA03-04, respectively. DNA fragments generated were assembled using NEBuilder HiFi DNA assembly kit (NEB). The resulting plasmid was verified by sequencing using primers oAA05-06.

### Bacillus directed mutagenesis

Steps leading to construction of new mutants are indicated in the last column of **Table 2** with additional details in this section. New mutants were often obtained by transformation of competent cells with a clean genomic DNA extracted from another existing *B. subtilis* mutants. Construction of some of the markerless class A PBPs mutants were previously described (12). Additional mutants described in this study were obtained using the plasmids pG+host9::*ΔponA,* pG+host10::*ΔpbpD,* pG+host10::*ΔpbpF* (**Table 1**) (12). PCR with primers oAG124 to oAG129 were consistently performed to check markerless deletions were not lost (12).

All transformations of the background strain with, and/or leading to, deletion of *ponA, mreB, mbl, gtaB, ugtP, ltaS* were selected for on NA medium supplemented with MgSO_4_ (20-25 mM) with the appropriate antibiotic.

To construct strain AG181 (*ΔmprF*), DNA fragments upstream and downstream the *mprF* coding sequence were amplified by PCR using primers oAG103-104 (2390 bp) and oAG105-106 (2414 bp), respectively and were digested by *Eco*RI and *Xba*I, respectively. The phleomycin resistance antibiotic cassette was amplified by PCR with primers PhleoXbaIfor and PhleoEcoRIrev from plasmid pIC22 and digested by both *Eco*RI and *Xba*I. The 3 digested PCR-fragments were ligated to generate a linear DNA, transformed into 168CA with selection for phleomycin resistance. The deletion of *mprF* was confirmed by PCR using primers oAG107-108.

Strain AG304 (*amyE::P_hyspank_mprF, spc*) was obtained by transformation of 168CA with pDR111-*P_hyspank_*-*mprF* plasmid (**Table 1**). Correct recombination was confirmed by the absence of amylase activity when grown on NA with 0.2% starch and by PCR (oAG300-301 primers). The same methods were used after transformation of other pDR111-derivatives.

We constructed our own *lytE* deletion mutant (*ΔlytE::cat*). For this, regions of genomic DNA upstream and downstream of *lytE* were amplified by PCRs with oAG350-351 (2048 bp) and oAG352-353 (2500 bp) primers pairs, respectively. These products were then digested by *Xba*I or *Bam*HI (**Table 3**). The chloramphenicol resistance cassette from pCotC-GFP plasmid (**Table 1**) was amplified with primers oAG197-198. The resulting PCR products (850 bp) was then digested by *Xba*I and *Bam*HI and the three DNA fragments were ligated and transformed into *B. subtilis* 168CA. Strain AG547 was obtained from this transformation and the deletion of *lytE* confirmed by PCR using primers oAG340-341 (**Table 3**).

Strains AG565 and AG586 were obtained by integration of pMUTin-‘*lytE-his* (erythromycin selection) in 168CA and *ΔcwlO* (AG474 strain) background, respectively. The plasmid integration (single cross-over) in *B. subtilis* resulted in expression of recombinant proteins with LEMRGSH_12_ tag in C-terminus, under the control of its native promoter. Since the integration of this plasmid into the *ΔcwlO* background did not result in any phenotypic change it indicated that the LytE-His_12_ was functional, as a double mutant *lytE cwlO* is lethal (44). Thereafter, genomic DNA of AG565 strain was used to transform all the strains needed for Western blot analysis (**Table 2**). Of note, AG1541 strain (*ΔcwlO*) obtained by genomic DNA recombination has the same genotype as AG586 strain obtained by plasmid integration.

### Random transposition mutageneses

Random transposition mutageneses were performed as previously described (39, 74) using the pMarB plasmid that carries the transposon TnYLB-1 encoding kanamycin resistance. pMarB was introduced into *B. subtilis* strains carrying pLOSS-*P_spac_*-*ponA* (12) (**Table 1**), a plasmid with an unstable origin of replication and *lacZ* which helps to monitor the plasmid stability by observing formation of blue or white colonies. After screening the random library of mutants, genomic DNA extracted from positive clones were backcrossed into the parental strain used for the transposon screen. Transposon insertion sites were identified by inverse PCR using gDNA extracted from the backcrossed positive clones (74) and sequencing the resulting PCR products. Further verification of the transposon’s site of insertion was obtained using specific primers. For the suppressor screen in *ponA pbpD pbpF* mutant background, we constructed strain AG141 (**Table 2**) which carries deletions of the three vegetative class A PBPs*, lacA* gene (74) and pLOSS-*P_spac_*-*ponA* (12). The transposon mutant library was screened on PAB agar supplemented with MgSO_4_ (10 mM), X-Gal and IPTG (1 mM) and white colonies selected. The strain SWA11a (**Table 2**) had a transposon in *mprF* coding sequence which was verified using specific primers oAG81-82 (**Table 3**).

A conditional lethal screen was also carried out in a *ponA pbpD pbpF mprF lacA* pLOSS-*P_spac_*-*ponA* strain (AG200, **Table 2**). Here the transposon mutant library was screened on NA and PAB supplemented with X-Gal and IPTG (1mM). Blue colonies indicating plasmid stability were selected. One mutant was isolated and its genomic DNA was backcrossed into AG200 background. The resulting blue forming AG200BK#42 strain (**Table 2**) was found to carry a transposon in *gtaB*’s coding sequence using primers oAG165-166 (**Table 3**).

### Spot growth assays

Strains were grown in NB with 20 mM MgSO_4_ at 30°C overnight. The cultures were then diluted 1:100 into fresh NB 10 mM MgSO_4_ (or when applicable 20 mM) and grown up to mid exponential phase at 37°C. Cells were harvested, washed in NB, and diluted to an OD_600nm_ ≈ 0.3 in NB. Each strain sample was then serially diluted 1:10 using a 96-multiwell plate and NB. Using a multichannel pipette, 5 μl of each dilution were transferred to various agar plates. As a control all the serial dilutions presented in the figures were also spotted on media supplemented with 10 mM or 20 mM MgS0_4_ to ensure that growth occurred for all spots (data not shown). Plates were incubated at 37°C overnight and scanned at different time points. Images were edited in Photoshop CS software.

### Microscopic imaging

Cells were grown to exponential growth phase at 37 °C in NB medium. Before the cells were mounted onto microscopic slides, the slides were covered with a thin layer of NA medium supplemented with the FM5-95 membrane dye at 140 μg/ml (Invitrogen). Fluorescence microscopy was carried out with a Zeiss Axiovert 200M microscope attached to a Sony Cool-Snap HQ cooled CCD camera with Nikon APO x100/1.40 oil Ph3 objective lens. Images were acquired using Metamorph 6 imaging software (Molecular Devices, Inc). Images were analysed with ImageJ (http://rsb.info.nih.gov/ij/) and assembled with Photoshop CS software. Images stained with the FM5-95 membrane dye were processed using the plugin ObjectJ in ImageJ (http://rsb.info.nih.gov/ij/) to measure cell diameter and cell length of about 400 cells per strain. Measurements were performed only once as cell diameter change was obvious under the microscope and on the images of **Fig 3D** (details in **Fig S3AB, Table 4**).

### Transmission electron microscopy

Strains were grown in NB medium (with IPTG when applicable) until reaching an OD_600nm_ of about 0.4. The cell pellets were fixed in 2% glutaraldehyde in sodium cacodylate buffer at 4°C for at least 24 h. Samples were spun and molten 4% agarose was added to the sample and re spun. Sample were allowed to cool in the fridge for 30 min. The agarose block was removed from each tube and cut into 1 mm^3^ pieces. The samples were then dehydrated by microwave processing using the Pelco Biowave® Pro+ incorporating the Pelco Coldspot® Pro. In the vacuum chamber, samples were rinsed 3 times in buffer (150 watts for 40s), then treated in 1% osmium, pulsed (100 watts for 8 min) and rinsed 3 times in distilled water (150 watts for 40s). They were then taken out of the vaccum chamber and a graded series of dried acetone was applied to the samples (25%, 50%,75%, and 3 times in 100%, at 150 watts for 40 s for each step). This was followed by impregnation with a series of increasing concentrations of epoxy resin (Taab Medium resin) in acetone (25%, 50%,75%, 3 times in 100% at 300 watts for 3 min. for each step). Samples were embedded in fresh 100% resin and polymerised at 60°C for 24 h in a conventional oven. Once polymerised the resin block were cut in semi-thin survey sections (0.5 μm) and stained with 1% toluidine blue in 1% borax. Ultrathin sections (70 nm approx.) were then cut using a diamond knife on a Leica EM UC7 ultramicrotome. The sections were stretched with chloroform to eliminate compression and mounted on Pioloform-filmed copper grids. Grids were stained with uranyl acetate (2%) and Lead citrate and viewed on a Hitachi HT7800 transmission electron microscope using an Emsis Xarosa camera.

### Samples preparation for LTA polymer detection

*B. subtilis* strains were grown overnight at 30°C with shaking in NB supplemented with 5 mM MgS0_4_ (or 10 mM for *mbl* mutants). The cells were then diluted (1:60-1:80 depending on the strain) to an approximate OD_600nm_ of 0.02 in prewarmed NB and NB supplemented with 0.2% glucose. Strains were then incubated at 37°C shaking and grown for about 4.5 h (late exponential phase). At the sampling time the OD_600nm_ of the cultures were measured before harvesting 15-25 ml of culture at 3273 *g* for 10 min, using a swing out centrifuge at room temperature. Pellets were washed with 200 μl of solution A (with 10 mL of Tris-HCl at 100 mM pH 7.4 and one cOmplete Mini EDTA-free tablet of protease inhibitor cocktail from Roche) and pelleted with benchtop centrifuge. Cells pellets were then frozen in liquid nitrogen and stored at −20°C until processing. Samples were resuspended in a volume of solution B (an equal volume of solution A and LDS sample buffer, Life Technologies) ranging from 70 to 200 μl in proportion to their OD_600nm_. Samples were boiled 30 min at 100°C, cooled down on ice for 3 min and incubated each with 100 U of benzonase (E1014, Sigma) for 30 min at 37°C. Samples were pelleted at 4°C, supernatants containing the extracted LTA were collected. Supernatants were frozen in liquid nitrogen and kept at −25°C. For the LTA Western blots, the quantity of sample loaded was normalized using the Bio-Rad Protein assays such that the final OD_595nm_ was approximately 0.35. Sample sets to be compared were grown, extracted, measured and normalized together for each experiment.

*S. aureus* strains were grown overnight at 37°C with shaking in LB. The cells were then diluted (about 300-fold) to an approximate OD_600nm_ of about 0.02 and grown in LB at 37°C until reaching late exponential and an OD_600nm_ of about 1. 10 ml of cultures were harvested, and samples were prepared as for *B. subtilis* with the following changes: 60 μl of solution B were used and about 18-20 μl of samples were loaded on SDS-PAGE with samples normalised to their OD_600nm_.

### Total protein samples preparation

For LytE or LtaS Western blots, cells were grown, harvested and processed in the same way as the LTA-samples, with the exception that to detect the His tagged proteins (LytE, LtaS or YfnI) the strains were grown in presence of erythromycin. An 8 ml culture sample was resuspended in 180-200 μl of a solution A and sonicated until a clear lysate was obtained. Samples were centrifuged at 16,000 *g* for 10 min at 4°C and a volume of sample supernatant loaded on the SDS-gel, normalized based on the OD_600nm_ of the cells culture collected.

### SDS-PAGE, LTA and proteins Western blots, detection blots

For SDS-PAGE, samples were diluted in solution B with the addition of reducing agent (1.5 μl per sample) in final volume of 20 μl, heated at 70°C for 10 min, loaded on the gel (NuPAGE 4-12% Bis-Tris gradient Midi Gel, Life Technologies) and separated by electrophoresis. For the LTA-PBP2B experiment, MES buffer, Novex Sharp prestained protein Ladder (Life Technologies) and 0.2 µm Amersham Hybond sequencing PVDF membrane (GE Healthcare) were used. Whereas for LytE (37 kDa) or LtaS (74 kDa) Western blots, MOPS buffer, Abcam prestained mid-range protein ladder (Ab116027) and 0.45 µM Amersham Hybond PVDF membrane (GE Healthcare) were used. The SDS-PAGE gels were transferred to membranes using the Trans-Blot Turbo transfer system (Bio-Rad) using an in-house transfer buffer (tris 600 mM, glycine 600 mM, tricine 280 mM, EDTA 2.5 mM and SDS 0.05%). After transfer, the membranes were washed 3x 5 min in PBS using a rocking shaker, afterwards the membranes were cut where required to allow different detection methods.

For the LTA Western blot, membrane was cut at the 60 kDa ladder position. The bottom part of the membrane was used for the LTA-blot whereas the top part was used to detect the membrane protein PBP2B (79 kDa) as a loading/extraction control. All LTA Western blot steps were performed at room temperature in a rolling shaker and using 50 mL falcon tubes. The LTA-membranes were incubated 1h15 in PBS buffer with 3% BSA (A7030, Sigma), then 1h30 in fresh PBS-3% BSA with a dilution 1:1000 of Gram-positive LTA monoclonal antibody (MA1-7402, Thermofischer). The membrane was then washed briefly 2 times, followed by 4x 8 min in PBS and then incubated in PBS-5% dried semi-skimmed milk with 1:10,000 anti-mouse HRP-linked secondary antibody (A9044, Sigma) for 1 h. This was then followed by 2 brief and 6x 8 min washes steps prior to detection.

For proteins Western blots to detect LytE-His_12_ or LtaS and PBP2B, all the following steps were performed at room temperature using a rocking shaker. The membranes were washed after transfer in PBS-Tween20 at 0.1% (here after noted PBS-T) and blocked in PBS-T with 5% milk for 1h15 (i.e PBS-T-milk) at room temperature. Thereafter the LytE-His_12_/PBP2B membrane blot was cut at about 53 kDa ladder position. Membranes were placed in fresh buffer PBS-T-milk for 1h (or 1h30 when performing the LTAs western blot in parallel) with the polyclonal rabbit anti-PBP2B antibody (at 1:5000, lab. stock or available at Merck ABS2199), the mouse monoclonal Penta-His antibody (Qiagen n°34660 stock at 200 µg/ml, used at dilution 1:2000) or polyclonal rabbit anti-LtaS antibody (at 1:1000, lab. stock). Two brief washing steps were performed in PBS-T followed by 3 washes of 10 min. Membranes were incubated in PBS-T-milk for 1 h using 1:10000 dilutions of anti-rabbit or anti-mouse HRP-linked secondary antibodies (A0545 or A9044 Sigma antibodies, respectively) as required. Membranes were then washed as described at the previous step.

Pierce ECL Plus Western Blotting Substrate (32132, Thermo Scientific) was used for detection as recommended by manufacturer. Chemiluminescence was detected using ImageQuant LAS 4000 mini digital imaging system (GE Healthcare) and membranes were exposed with 1 min increment for 20 min. Membranes were also imaged by Epi-illumination to detect the pre-stained proteins ladder on the membrane. In our conditions, quantities of LTAs samples loaded were scaled down to avoid burn out of signals observed in some of the tested strains. Occasionally for the LTAs blot, strains (in NB-only) can display a very weak signal above the 30 kDa ladder position. This phenomenon was observed by others performing *Bacillus* LTA Western blots and was assumed not to be related to LTA but possibly a cross-reactivity of the antibody to WTAs that also contains glycerolphosphate (18).

Images were edited in ImageJ (Fiji), the same contrast enhancement was applied for the membranes of the same set experiment. Three independent sets of experimental samples were generated by Western blots. Contrast was sometimes added to the images, in particular to observe LytE signal in low-expressing strains.

### LytE purification and raising antiserum

Plasmid pAM-21 was transferred into *E. coli* BL21 (DE3) strain. Newly obtained strain grew at 37°C in LB kanamycin, when cell reached an approximative OD_600nm_ of 0.5 *lytE* expression was induced with 1 mM IPTG for 3 h. 50 ml culture was collected by centrifugation at 9,000 *g* for 5 min and resuspended in 15 ml of a solution of PBS at 4 µg/ml of lysosyme. After incubation for 20 min at room temperature (RT), cells were sonicated, and once clear lysate was obtained cells were harvested at 3,000 *g* for 4 min. The cell pellet was resuspended in PBS buffer at 8 M Urea and incubated for 30 min at RT on a rolling shaker. The cells were then centrifuged at 9,000 *g* for 10 min and supernatant was collected and mixed gently with 200 µl of Ni Sepharose High Performance (GE Healthcare) and incubated for 1 h in same condition as above. Then the sample was washed 3 to 4 times in buffer A (50 mM Tris (pH 8) and 300 mM NaCl), and the last wash was done in buffer B (50 mM Tris, 300 mM NaCl, 5 mM Imidazole). Sample was eluted five times in buffer C (PBS at 8 M Urea and 100 mM imidazole) and elution fractions were analysed by SDS-PAGE. Ice-cold acetone was added at a 4:1 volume ratio to the desired eluted sample and incubated overnight at −20°C. Sample was centrifuged at 4 °C for 10 min at 15,000 *g* and the pellet was left to dry at RT. The pellet was suspended in 200 µl of PBS and 100 µg aliquot was sent to raise antiserum (Eurogentec). The LytE antiserum was used in Western blot at 1:5000 dilution and with the anti-rabbit secondary antibody (1:10000).

### Growth assays in presence of Daptomycin

For the *S. aureus* samples for LTA Western blot, cultures were done as previously described but after 3h growing in LB, CaCl_2_ (1.25 mM final) was added to the cultures which were then divided and grown an extra hour with or without daptomycin (Dap stock at 1 mg/ml, Abcam ab141204).

For the NA-CaCl_2_ DAP assays, strains were exponentially growing in NB for about 3h at 37°C (exponential phase) and then normalized to an OD_600nm_ of 0.5. Cells (300 µl) were spread on calibrated NA plate supplemented with CaCl_2_ (1.25 mM final). A daptomycin 0.016-256 mg/l antimicrobial susceptibility testing strip (Liofilchem) was applied to the medium surface. Plates were incubated at 37°C overnight and scanned after 24h. Images were edited in Fiji (ImageJ). Interpretation on growth inhibition follows manufacturer’s guidance.

To monitor strain growth in a plate reader, we used pre-cultures which has grown in NB (or in LB for *S. aureus*) at 37°C for about 3h. For each assays, a daptomycin stock at 17.4 mg/ml was used to prepare a fresh DAP stock at 0.1 mg/ml and this was used to make dilutions of the antibiotic as required in 96-wells plates in 100 µl of medium supplemented with CaCl_2_ (1.25mM final). Plates were then loaded with 100 µl of cells (diluted in medium-CaCl_2_ and normalized at the desired OD_600nm_). Plates were shaken at 37°C in a Tecan Sunrise plate reader and the OD_600nm_ was measured every 6 min. Conditions were tested in duplicate or triplicates whenever possible. Data were collected using Magellan software (version 7.2). Microsoft Excel was used for analysis and generating graphs.

## Acknowledgements

The authors thank P. Aldridge for critical reading of the manuscript. The authors thank colleagues for strains, genomic DNAs or daptomycin stock: S. Moore, P. Dominguez-Cuevas, P. Gamba, A. Koh, K. Seistrup, L. Bowman, J. Buttress, H. Strahl and J. D. Helmann, B. M. Wendel and A. Peschel for strains related to daptomycin resistance. Technical support was provided by I. Selmes, F. Davison and T. Fletcher and TEM sample processing by Tracey Davey in the Electron Microscopy Research Services Newcastle University (funded through the BBSRC; BB/R013942/1). The authors thank J. Errington for the anti-LtaS antibody, H. Strahl and W. Vollmer for helpful discussion, C. Winterhalter and S. Fenyk for advice on Western blots and P. Tiago for sharing subtiwiki data. This work was funded by grants from the BBSRC (BB/G015902/1), EPSRC (EP/N031962/1) (RAD and AG) and SACB (studentship for AA).

## Author contributions

Conceptualization: AG; Funding acquisition, RAD; Methodology: AG; Investigation: AG and AA; Methodology: AG; Project administration: RAD; Supervision: RAD; Validation: AG; Visualization: AG; Writing—original draft: AG; Writing—review & editing: AG and RAD.

## Supporting information

Figures, Tables and Supplemental Figures are available for this manuscript. This study includes no data deposited in external repositories.

