## supplemental figures for "Insights into the role of lipoteichoic acids and MprF function in *Bacillus subtilis*"

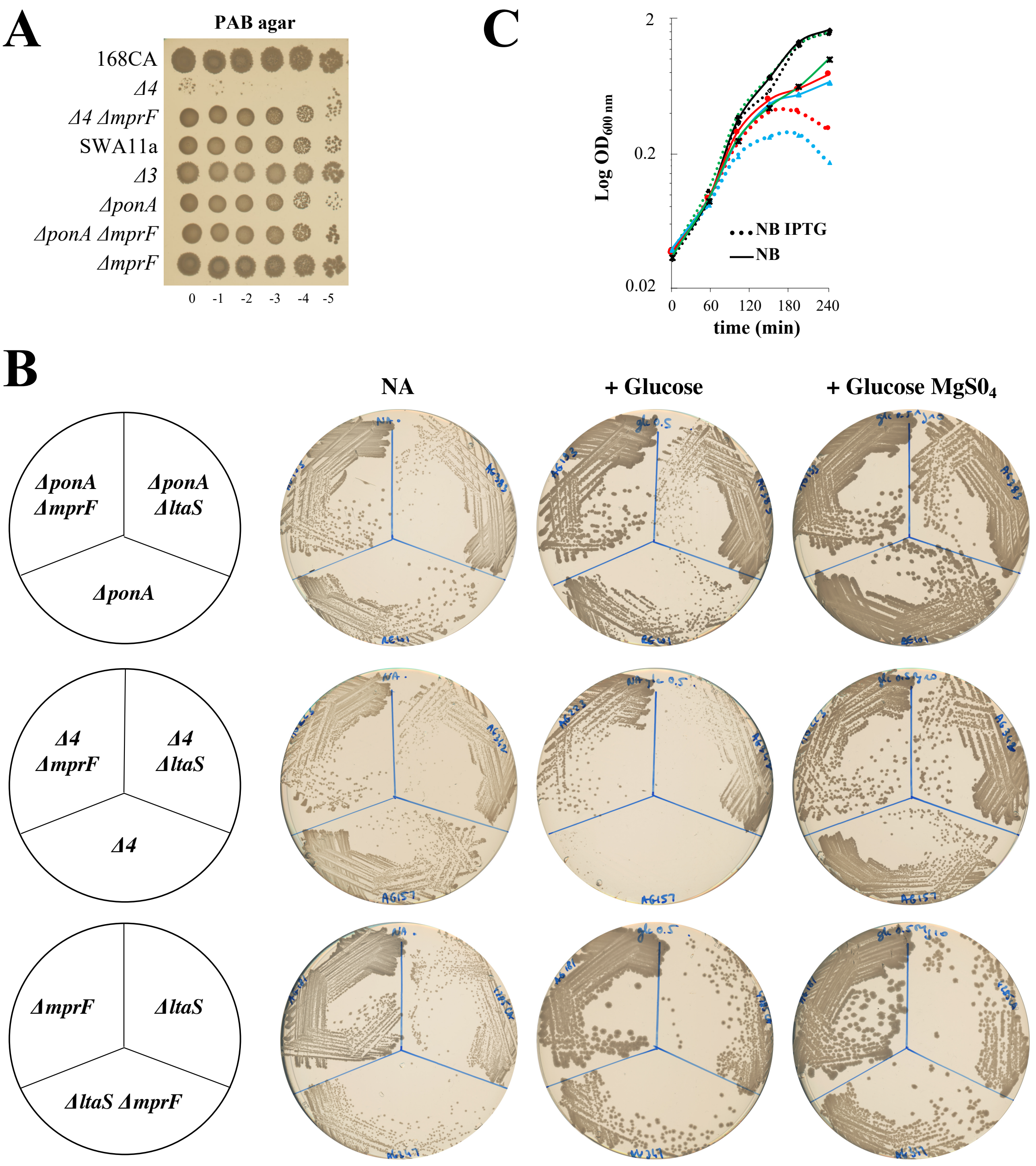


**Figure S1. Conditional growth of mutants in cell envelope components.**

**(A)** Ten-fold spot dilution growth assays of strains *B. subtilis* wild type 168CA, AG157 (*ΔponA ΔpbpD ΔpbpF ΔpbpG* renamed *Δ4*), AG223 (*Δ4 ΔmprF*), SWA11a (*ΔponA ΔpbpD ΔpbpF mprF::TnYLB-1 lacA*), AG417 (*ΔpbpD ΔpbpF ΔpbpG* renamed *Δ3*), RE101 (*ΔponA*), AG193 (*ΔponA ΔmprF)* and AG181 (*ΔmprF*). Serial dilutions were spotted onto nutrient agar (NA) with or without glucose 1% (**Fig 2A**) as well as on PAB agar (this figure). Plates were incubated at 37°C for 24 h and imaged.

**(B) Comparison of *Δ4* and *ΔponA* mutants in absence MprF or LtaS.** Strains were streaked from glycerol stocks onto NA plates (supplemented with antibiotic and/or magnesium, where required). After incubation at 37°C overnight, a single colony for each strain was then streak across the set of plates shown above (with glucose 0.5%, $MgS0_4$ 10mM) and scanned after 16 h incubation at 37°C. The following strains were inoculated: RE101 (*ΔponA*), AG193 (*ΔponA ΔmprF*), AG383 (*ΔponA ΔltaS*), AG157 (*Δ4*), AG223 (*Δ4 ΔmprF*), AG342 (*Δ4 ΔltaS*), AG181 (*ΔmprF*), 4285CA (*ΔltaS*), AG347 (*ΔmprF ΔltaS*).

**(C)** Overnight cultures of strains were diluted back in NB and grown at 37°C in presence or absence of IPTG (dashed or solid lines, respectively). Growth curves of strains carrying $P_{hyspank}$-*mprF* are displayed with the following code: wild-type-like MprF ↑ (black, diamond symbol, AG304 strain), *ΔponA* MprF ↑ (red, round symbol, AG311 strain), *Δ4* MprF ↑ (blue, triangle symbol, AG317 strain) and *Δmbl* MprF ↑ (green, square-cross symbol, AG322 strain).

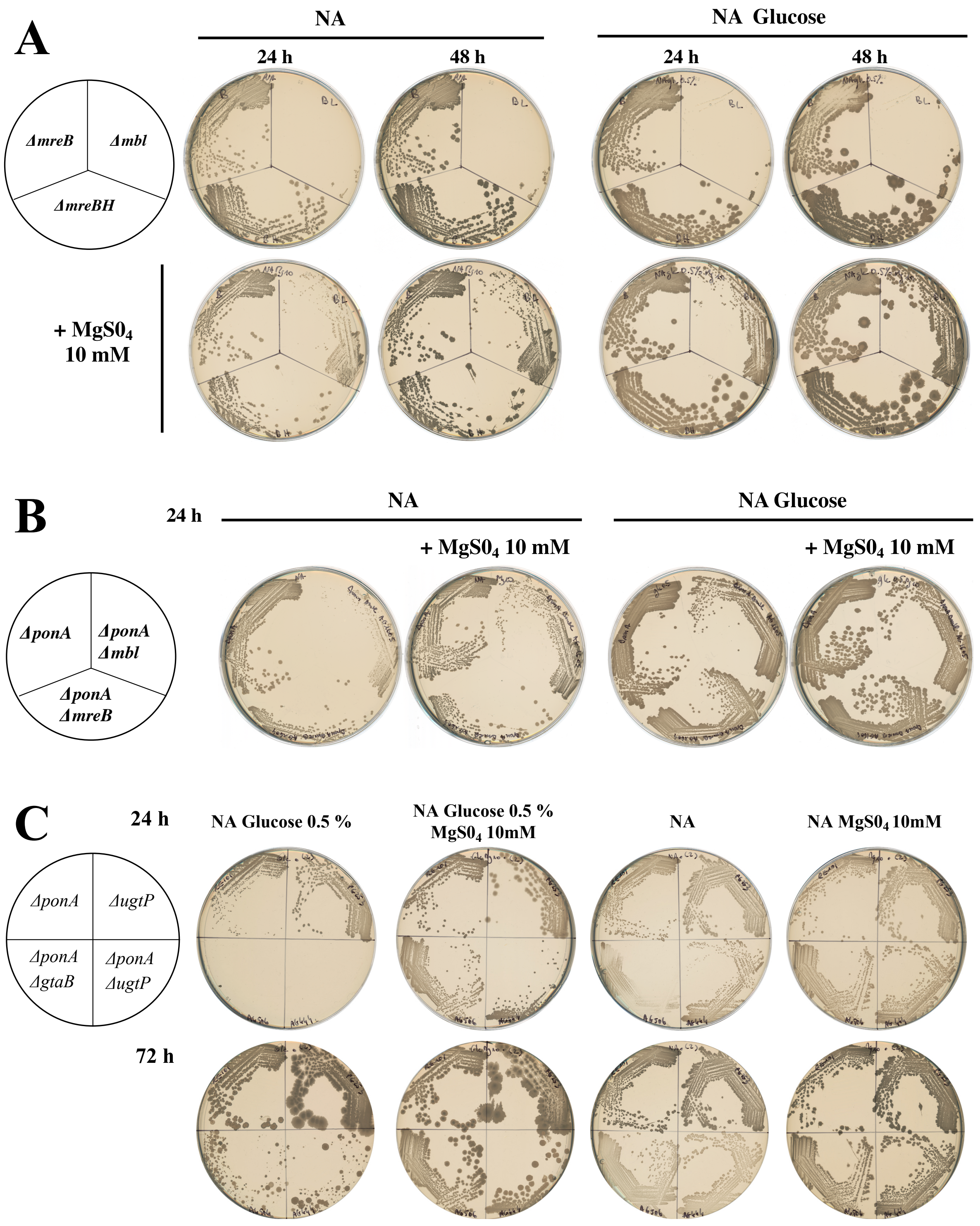


**Figure S2. Conditional growth of strains with magnesium dependence.**

**(A-B) Growth of actin like mutants in 168CA background and in absence of *ponA*.** Strains were streaked from the glycerol stock onto NA plates supplemented with antibiotic and/or magnesium, where required. After incubation at 37°C overnight, a single colony for each strain was then streak across the set of plates (glucose 0.5%) shown above and scanned after 24 h or 48 h incubation at 37°C. The following strains obtained in 168CA background were inoculated: (**A**) *ΔmreB* (KS36), *Δmbl* (AK045B), *ΔmreBH* (AG1593); (**B**) *ΔponA* (RE101), *ΔponA Δmbl* (AG1605), *ΔponA ΔmreB* (AG1604).

(**C**) **Absence of *gtaB* or *ugtP* causes severe growth defect in absence of *ponA*.** Strains *ΔponA* (RE101), *ΔugtP* (PG253), *ΔponA ΔugtP* (AG444), *ΔponA ΔgtaB* (AG506) were prepared as described for A-B and were then streaked across a set of plates shown above. Pictures were taken after 24 h and 72 h incubation at 37°C.

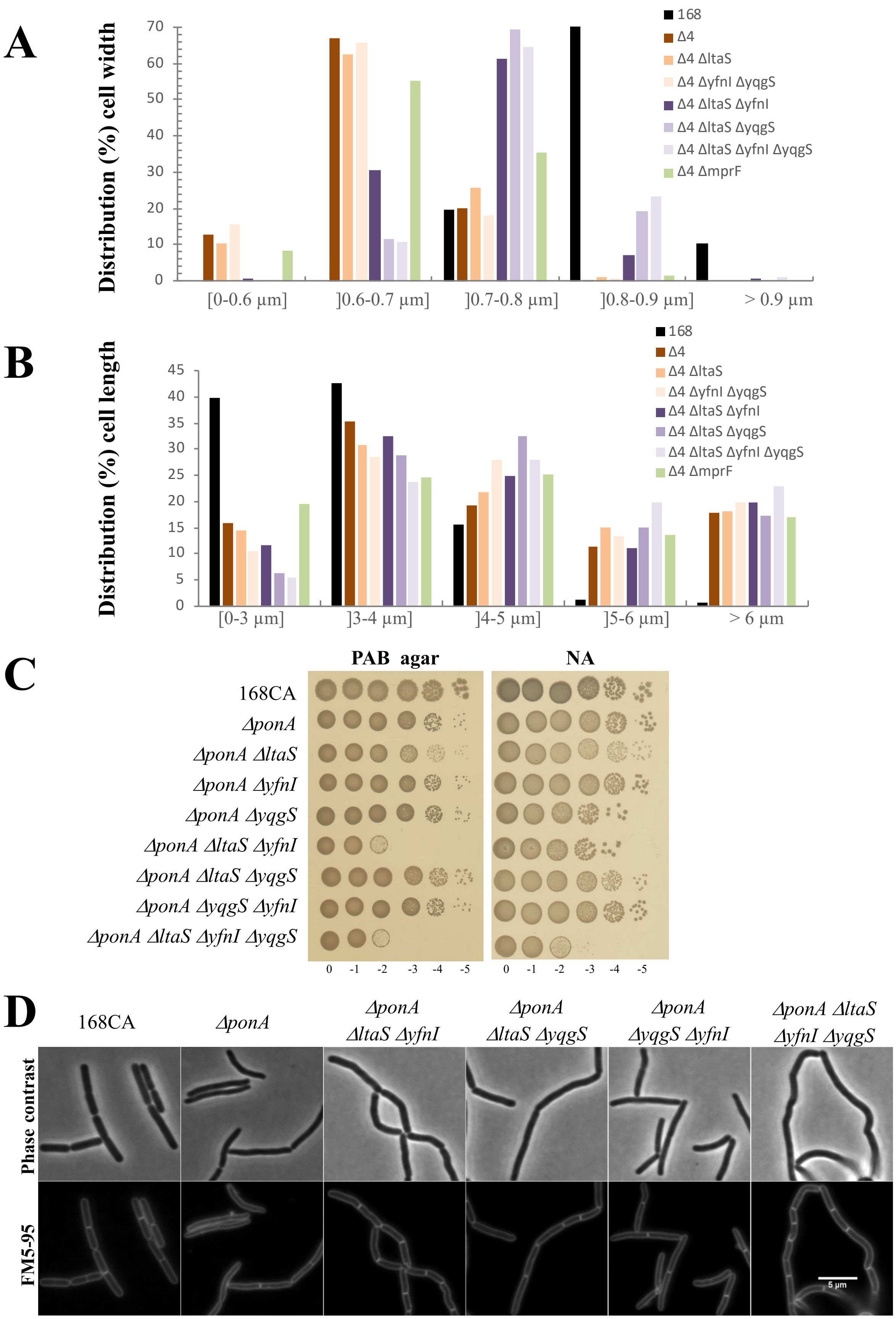


**Figure S3.** **Cell morphology changes associated with deletion of LTA-synthase genes combined with class A PBPs null mutants of *B. subtilis.***

**(A-B)** All the strains grew exponentially in NB with 10 mM $MgS0_4$, were washed and diluted in NB. Strains 168CA, AG157 (*Δ4*), AG342 (*Δ4 ΔltaS*), AG377 (*Δ4 ΔyfnI ΔyqgS*), AG370 (*Δ4 ΔltaS ΔyfnI*), AG372 (*Δ4 ΔltaS ΔyqgS*), AG380 (*Δ4 ΔltaS ΔyfnI ΔyqgS*) and AG223 (*Δ4 ΔmprF*) (**Table 2**) grown for 120 min until mid-exponential at 37°C. Cells were stained with FM5-95 dye and microscopy images were acquired. Cells were measured using plugin ObjectJ in ImageJ. About 400 cells were counted for each strain. Content of the figure is from data acquired by one experiment only. Due to the formation of twisted cell chains, the cells measurement of *Δ4 ΔltaS ΔyfnI ΔyqgS* strain was done on the straight regions of the chain. Distributions of cell width **(A)** and cell length **(B)**. **Table 4** indicates the average of cell width and cell length in the cell population analysed.

**(C-D). Effect of LTA-synthase deletions in *ΔponA* background. (C)** 10-fold serial dilutions were prepared in NB as in previous assays for strains 168CA, RE101 (*ΔponA*), AG383 (*ΔponA ΔltaS*), AG384 (*ΔponA ΔyfnI*), AG385 (*ΔponA ΔyqgS*), AG393 (*ΔponA ΔltaS ΔyfnI*), AG394 (*ΔponA ΔltaS ΔyqgS*), AG395 (*ΔponA ΔyfnI ΔyqgS*) and AG403 (*ΔponA ΔltaS ΔyfnI ΔyqgS*) (**Table 2**). Plates were incubated at 37°C and imaged at 22 h. (**D)** Strains described in (**C**) were diluted in NB and grown at 37°C for 120 min. When cells reached mid-exponential phase, cell membranes were stained with a FM5-95 dye and observed under a microscope. Representative images were assembled in photoshop. Scale bar represents 5 µm.

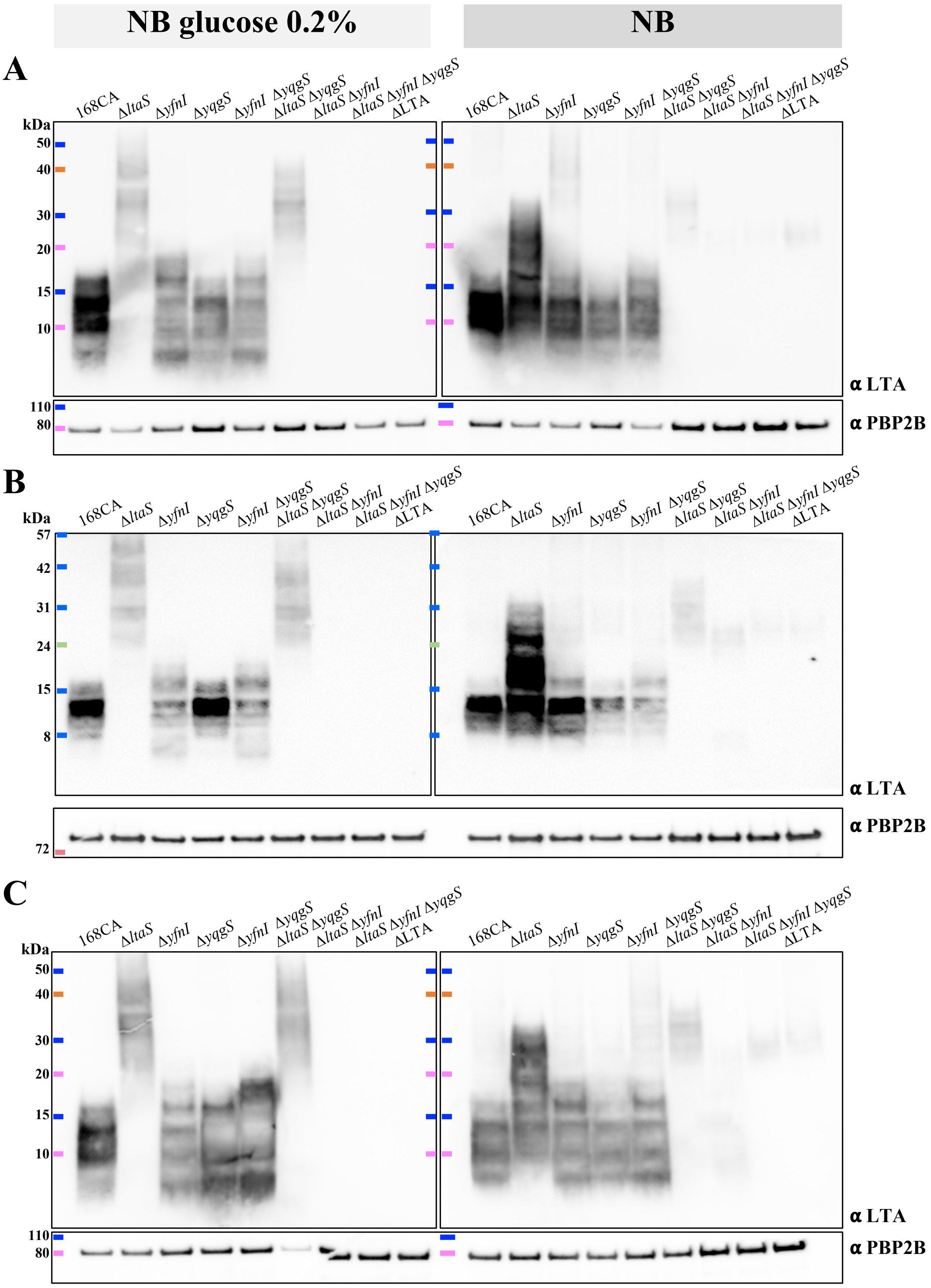


**Figure S4. Results of three independent LTA Western blots supporting Fig 4A.**

The lower part of the membrane was probed using a monoclonal LTA antibody and an HRP-linked anti-mouse antibody. The other membrane part was probed using polyclonal anti-PBP2B and HRP-linked anti-rabbit antibodies, to provide a sample loading control. Samples were prepared from strains *B. subtilis* wild type (168CA), 4285CA (*ΔltaS*), 4289CA (*ΔyfnI*), 4292CA (*ΔyqgS*), AG595 (*ΔyfnI ΔyqgS*), AG593 (*ΔltaS ΔyqgS*), AG594 (*ΔltaS ΔyfnI*), AG600 (*ΔltaS ΔyfnI ΔyqgS*) and 4620 (*ΔltaS ΔyfnI ΔyqgS ΔyvgJ*, noted here *Δ*LTA).

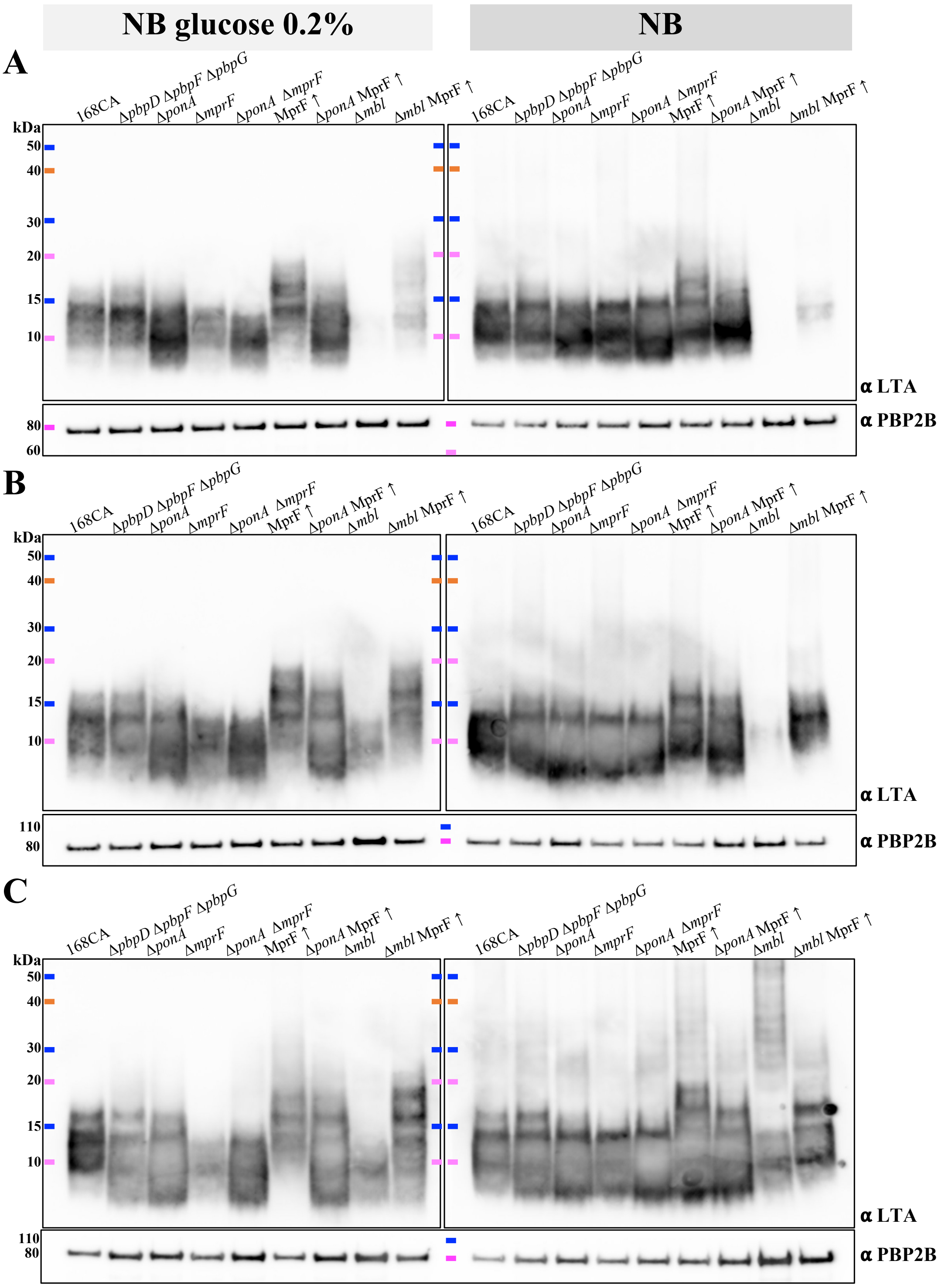


**Figure S5. Results of three independent LTA Western blots supporting Fig 4B.** ***To continue***

**D**

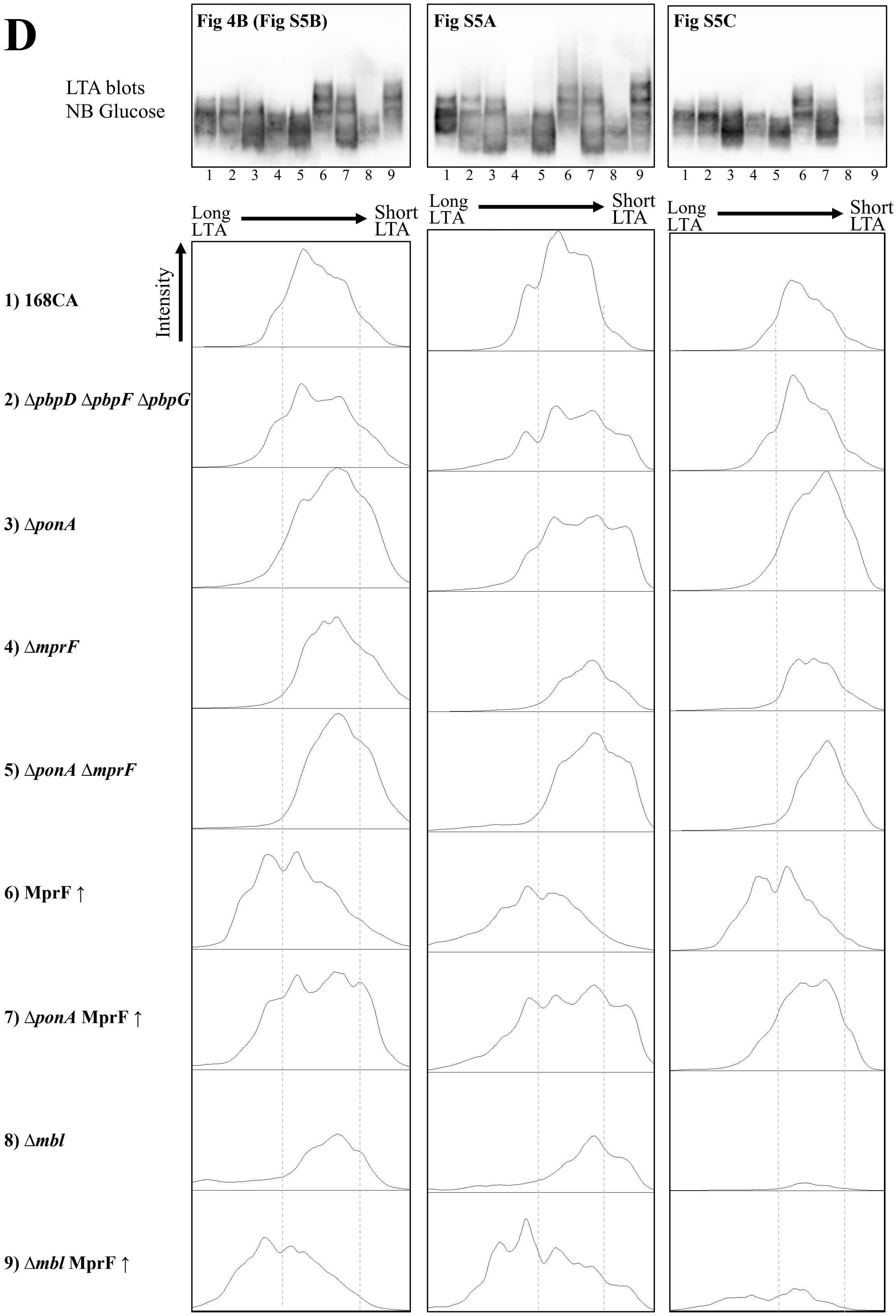


60 **Figure S5. Results of three independent LTA Western blots supporting Fig 4B.**

(**A-C**)The lower part of the membrane was probed using a monoclonal LTA antibody and an HRP-linked anti-mouse antibody. The other membrane part was probed using polyclonal anti-PBP2B and HRP-linked anti-rabbit antibodies, to provide a sample loading control. Samples were prepared from strains *B. subtilis* wild type (168CA), AG417 (*ΔpbpD ΔpbpF ΔpbpG*), RE101 (*ΔponA*), AG181 (*ΔmprF*), AG193 (*ΔponA ΔmprF*), AG304 (MprF ↑), AG311 (*ΔponA* MprF ↑), 4261CA (*Δmbl*), AG322 (*Δmbl* MprF ↑).

(**D**) The LTA signals of some of the strains grown with glucose are represented here as densitometry graphs below their corresponding LTA blot. Analysis was done for each biological replicate presented in either **Fig 4B (=Fig S5B), Fig S5A, Fig S5C.** Raw blot images were analysed using ImageJ software and the function ‘analysis gels’ after a same size area was defined for each lane. The dashed lines provide guidelines to help compare the shift of LTA length between samples relative to the wild type.

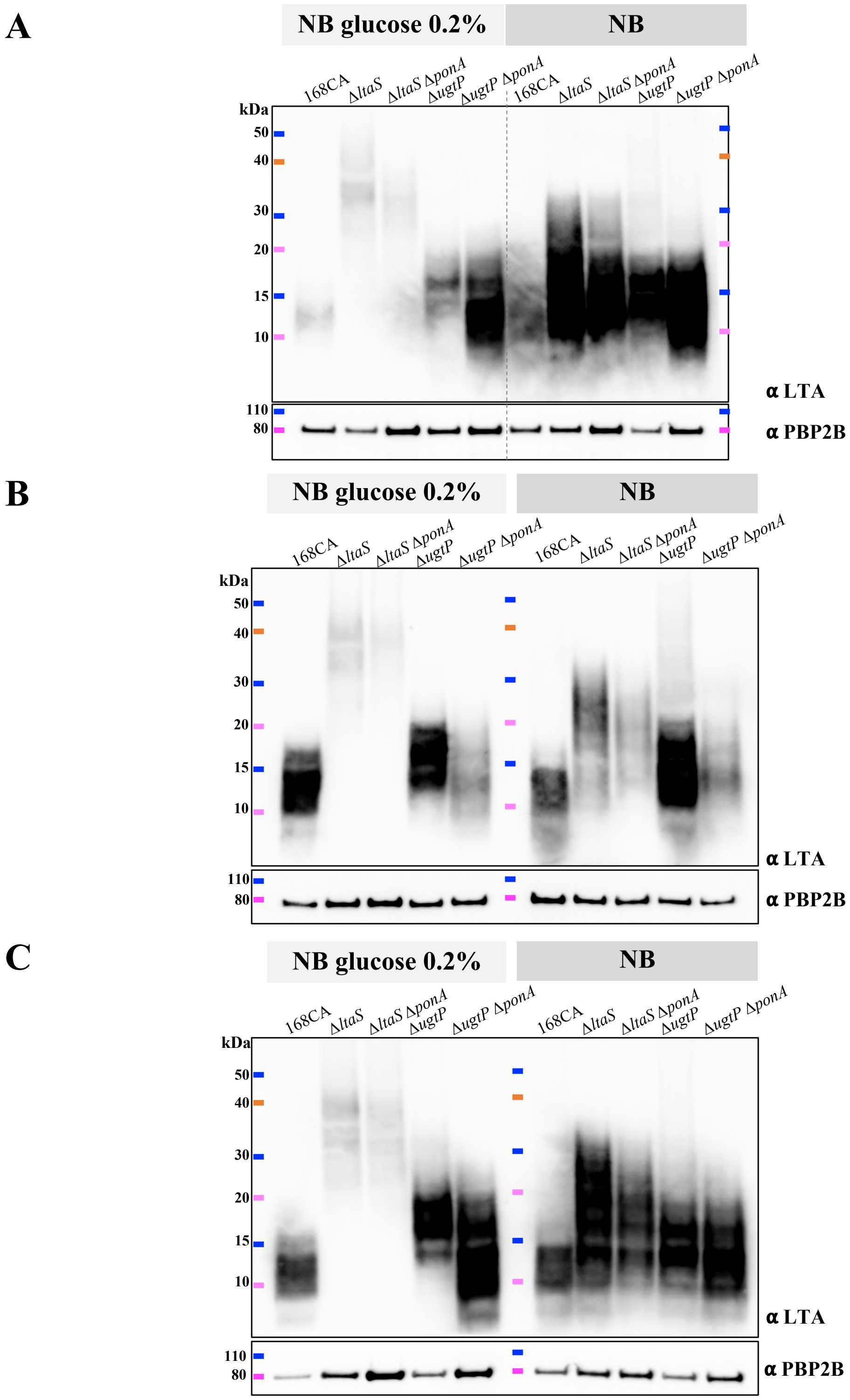


**Figure S6. Results of three independent LTA Western blots supporting Fig 4C.** The lower part of the membrane was probed using a monoclonal LTA antibody and an HRP-linked anti-mouse antibody. The other membrane part was probed using polyclonal anti-PBP2B and HRP-linked anti-rabbit antibodies, to provide a sample loading control. Samples were extracted from *B. subtilis* 168CA, 4285CA (*ΔltaS*), AG383 (*ΔltaS ΔponA*), PG253 (*ΔugtP*) and AG444 (*ΔugtP ΔponA*) strains.

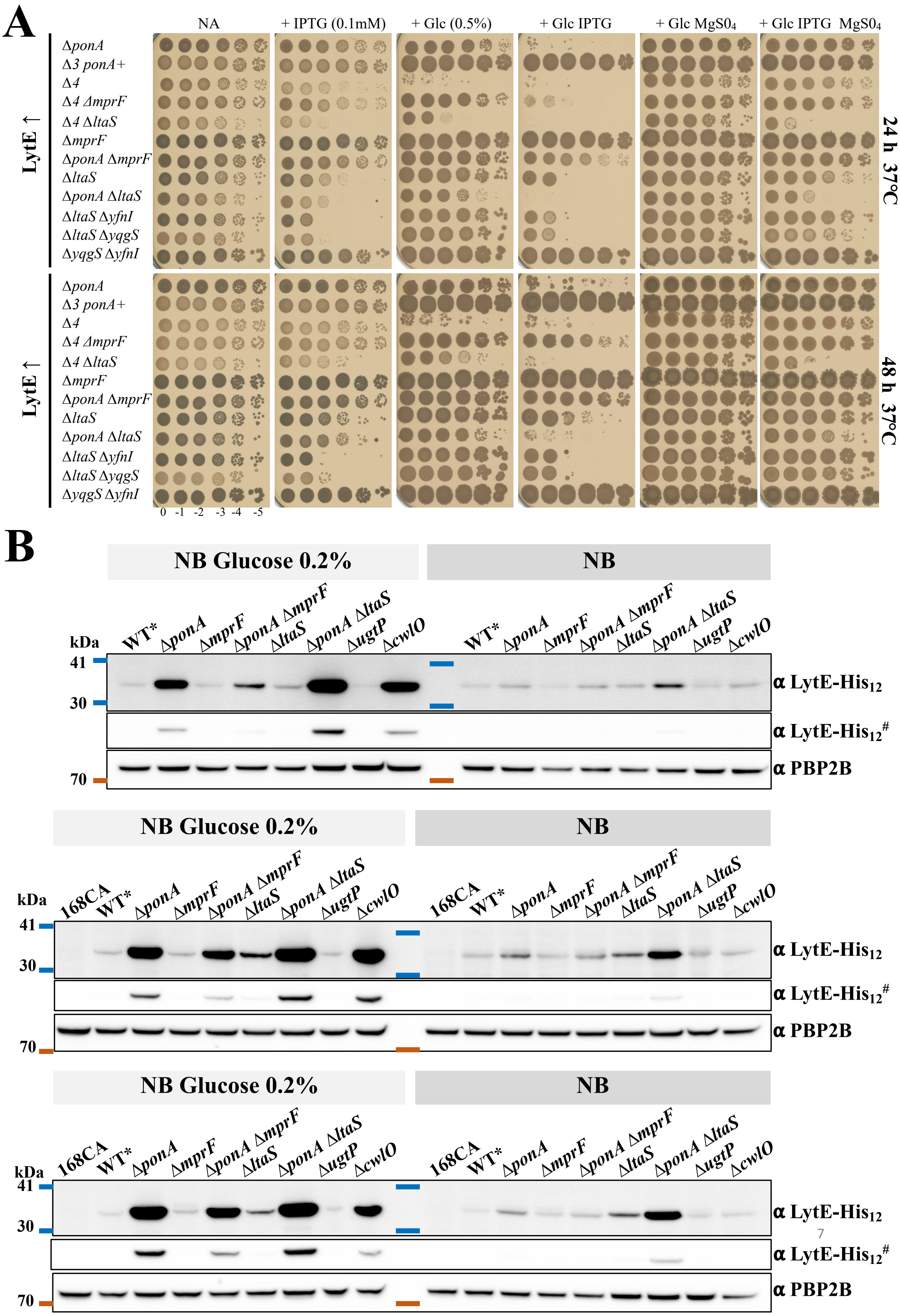


79 **Figure S7**: **Abundance of LytE in the cell envelope and strains growth characteristics.**

**(A) LytE overexpression is conditionally lethal in absence of PBP1 or LtaS.** *lytE* overexpression was analysed in strains which integrated at the *amyE* locus an IPTG-inducible $P_{hyspank}$-*lytE,* represented here LytE ↑. The strains tested had the following relevant features: AG484 (*ΔponA* LytE ↑), AG1460 (*Δ3 ponA*$^{+}$ LytE ↑), AG501 (*Δ4* LytE ↑), AG502 (*Δ4 ΔmprF* LytE ↑), AG1465 (*Δ4 ΔltaS* LytE ↑), AG1478 (*ΔmprF* LytE ↑), AG1479 (*ΔponA ΔmprF* LytE ↑), AG1462 (*ΔltaS* LytE ↑), AG1480 (*ΔponA ΔltaS* LytE ↑), AG1497 (*ΔltaS ΔyfnI* LytE ↑), AG1498 (*ΔltaS ΔyqgS* LytE ↑) and AG1499 (*ΔyqgS ΔyfnI* LytE ↑). The following strains that did not display any obvious phenotype are not represented here: AG475 (168CA LytE ↑), AG1461 (*ΔlytE* LytE ↑), AG1463 (*ΔyfnI* LytE ↑), AG1492 (*ΔyqgS* LytE ↑). NA plates were supplemented with IPTG (0.1 mM), glucose Glc (0.5%) and $MgS0_4$ (10 mM). Plates were incubated at 37°C and scanned after 24 h (**Fig 5A**) and 48 h.

**(B) Results of three independent experiments supporting Fig 5C.** Relative abundance of LytE and PBP2B protein in mutant strains detected by Western blots. Strains expressing LytE-His$_{12}$ under the control of its native promoter were grown in NB and NB supplemented with glucose 0.2% until late exponential growth at 37°C. For simplification only the relevant background features are displayed for the following strains: wild type-like AG565 (WT*), AG587 (*ΔponA*), AG1486 (*ΔmprF*), AG1487 (*ΔponA ΔmprF*), AG1489 (Δ*ltaS*), AG1488 (*ΔponA ΔltaS*), AG1535 (*ΔugtP*) and AG1541 (*ΔcwlO*). In addition, the wild type 168CA (i.e. not expressing LytE-His$_{12}$) was grown in parallel and used here as a negative control for the LytE-blot. The bottom part of the membrane (split at ~ 53 kDa position) was incubated with a monoclonal Penta-His and an HRP-linked anti-mouse antibodies. The top membrane part was detected with a polyclonal anti-PBP2B and HRP-linked anti-rabbit antibodies. # For each section, the middle panel corresponds to the raw image of the LytE-His$_{12}$ blot at a detection time where one of the sample signals had reached saturation.

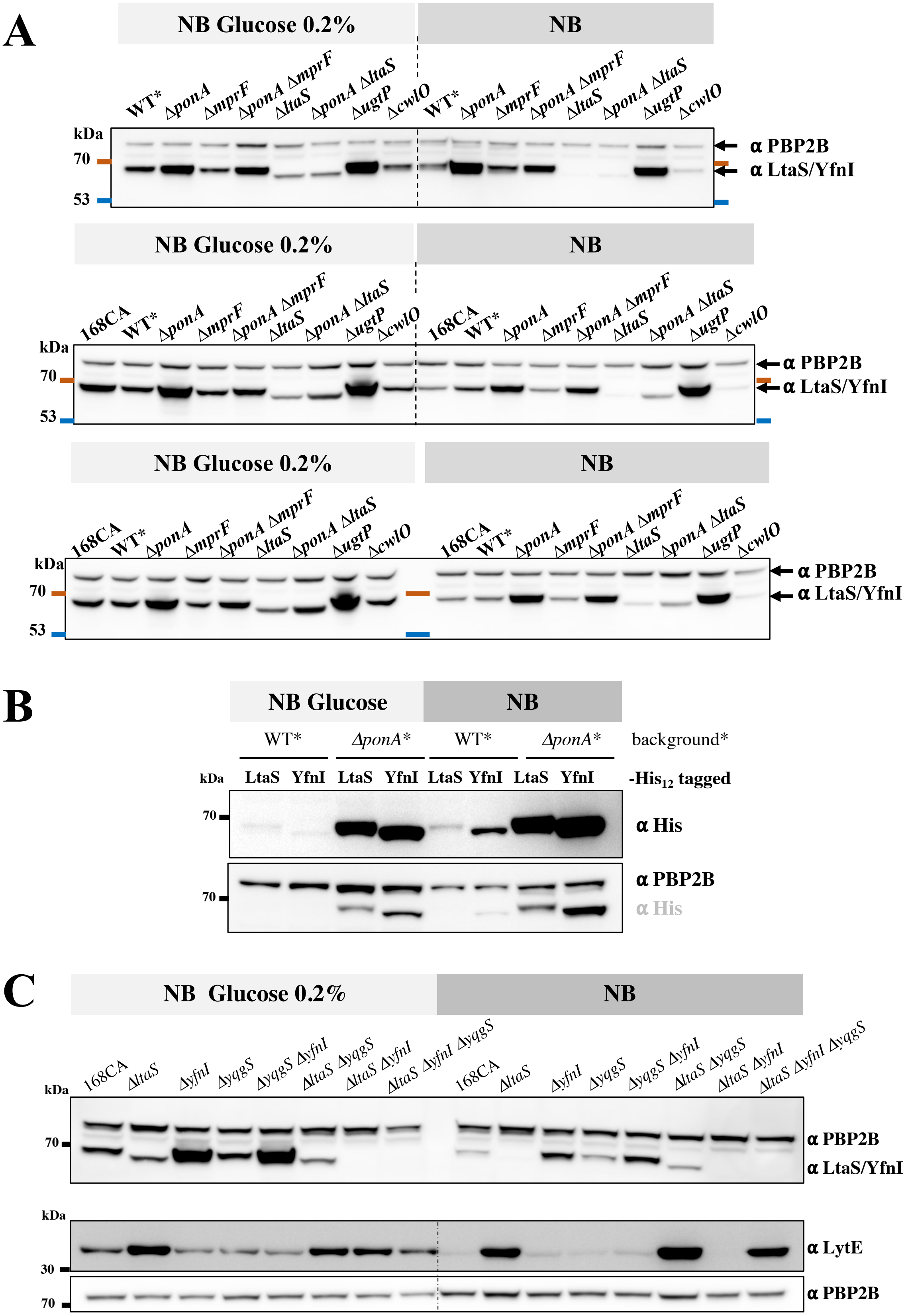


**Figure S8**. **Data supporting the results of Fig 5.**

**(A) Results of three independent experiments supporting Fig 5D**. Relative abundance of LTA-synthases LtaS, YfnI and PBP2B protein levels in mutant strains detected by Western blots. The samples used in **Fig S7B** were loaded on a new SDS-gel and the Western blot was developed with polyclonal anti-LtaS (that cross reacts with YfnI, unpublished Errington's lab), anti-PBP2B antibodies, and HRP-linked anti-rabbit antibody. A weak non-specific protein band is observed between PBP2B and LtaS and is detected by the anti-PBP2B antibody. Strains expressing LytE-His$_{12}$ under the control of its native promoter were grown in NB and NB supplemented with glucose (0.2%) until late exponential phase at 37°C. For simplification only the relevant background feature is displayed for the following strains: 168CA, wild type-like AG565 (WT*), AG587 (*ΔponA*), AG1486 (*ΔmprF*), AG1487 (*ΔponA ΔmprF*), AG1489 (Δ*ltaS*), AG1488 (*ΔponA ΔltaS*), AG1535 (*ΔugtP*) and AG1541 (*ΔcwlO*).

**(B) Accumulation of LtaS-His and YfnI-His in *ΔponA* cells.** Strains were grown in NB and NB supplemented with glucose 0.2% until late exponential growth at 37°C. Tested once in condition similar to that of the proteins Western blot presented in this study, following a preliminary test. * Strains used here express the following his tagged protein under the control of their native promoter: LtaS-His$_{12}$ in a wild type-like background (AG569) or in *ΔponA* (AG588) background; YfnI-His$_{12}$ in a wild type-like (WT) (AG570 strain) or in *ΔponA* (AG589) background. His tagged proteins were detected with a monoclonal Penta-His and an HRP-linked anti-mouse antibodies. The membrane was reused to detected PBP2B (sample loading control) with a polyclonal anti-PBP2B and HRP-linked anti-rabbit antibodies (second panel). Experiment tested on one biological set of samples.

**(C) Accumulation of LtaS/YfnI and LytE in LTA-synthase mutants.** The following strains were grown (NB ± Glucose 0.2%) in condition similar to our previous assays: *B. subtilis* 168CA, 4285CA (*ΔltaS*), 4289CA (*ΔyfnI*), 4292CA (*ΔyqgS*), AG595 (*ΔyqgS ΔyfnI*), AG593 (*ΔltaS ΔyqgS)*, AG594 (*ΔltaS ΔyfnI)*, AG600 (*ΔltaS ΔyfnI ΔyqgS*). In this figure, the Western blot experiments were carried out in same condition as other assays (polyclonal anti-LtaS antibody cross reacts with YfnI, unpublished Errington's lab) (**A**), except for the use of our newly produced polyclonal anti-LytE antibody (**C**). Experiment tested on one biological set of samples.

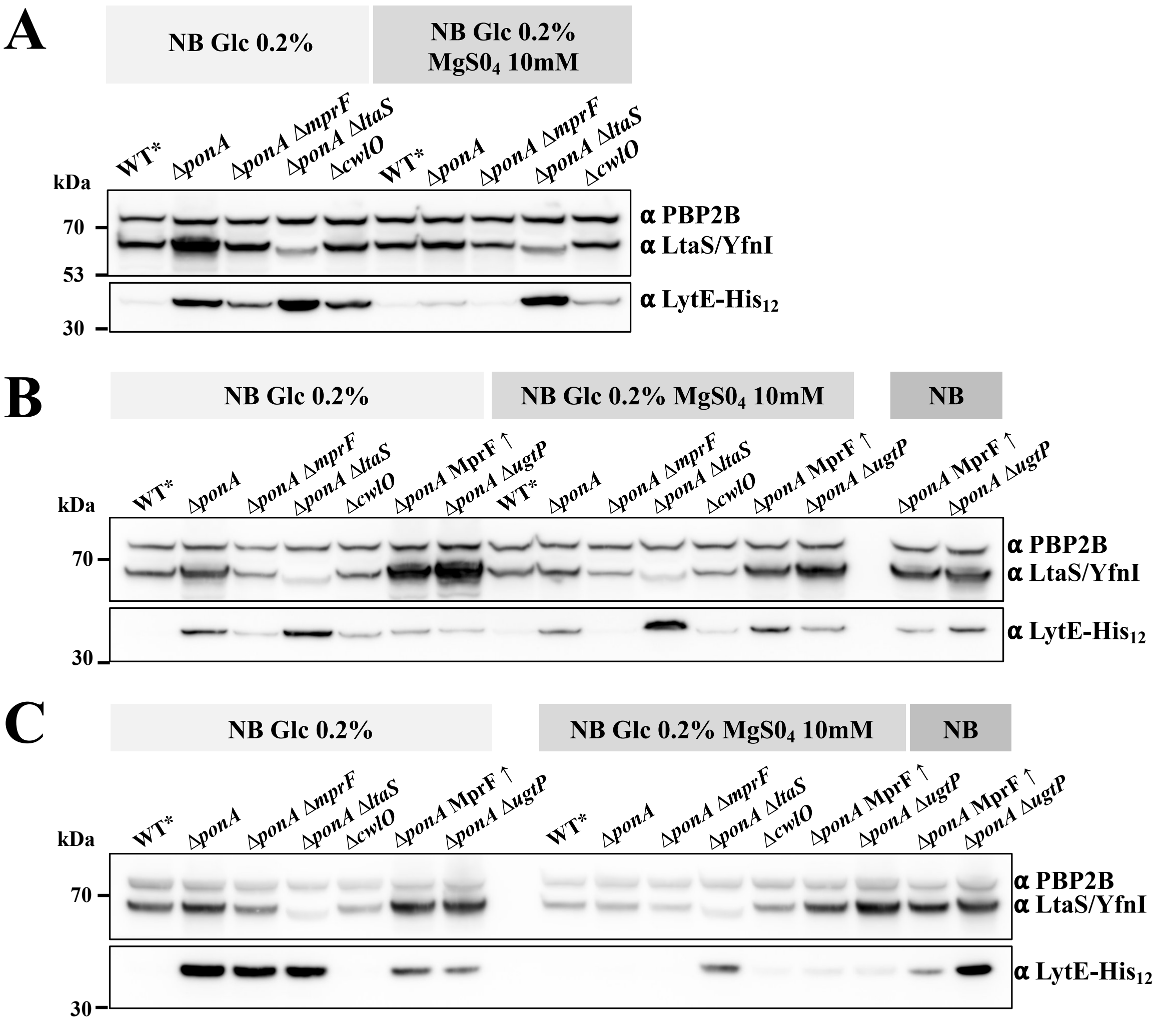


**Figure S9. Effect of magnesium in culture medium on the production of LytE and LtaS and accumulation of LtaS/YfnI and LytE in synthetically sick strains Δ*ponA* MprF ↑ and Δ*ponA* Δ*ugtP*.**

Detection of LytE-His$_{12}$ and PBP2B by Western blots. Strains expressing LytE-His$_{12}$ under the control of its native promoter were grown in NB supplemented with glucose 0.2% with or without MgS0$_4$ (10 mM) at 37°C. For simplification on this figure only the relevant background feature is displayed for the following strains: wild type like AG565 (WT*), AG587 (*ΔponA*), AG1487 (*ΔponA ΔmprF*), AG1488 (*ΔponA ΔltaS*), AG1541 (*ΔcwlO*), AG1684 (*ΔponA* MprF ↑) and AG1685 (*ΔponA ΔugtP*). The last two strains were also grown in parallel in NB. The top membrane part was incubated with polyclonal anti-LtaS (that cross reacts with YfnI, unpublished Errington's lab), anti-PBP2B antibodies, and HRP-linked anti-rabbit antibody. The bottom part of the membrane was incubated with a monoclonal Penta-His and an HRP-linked anti-mouse antibodies. Images were processed in ImageJ software. Results of three independent experiments (**A-C**) are presented here.

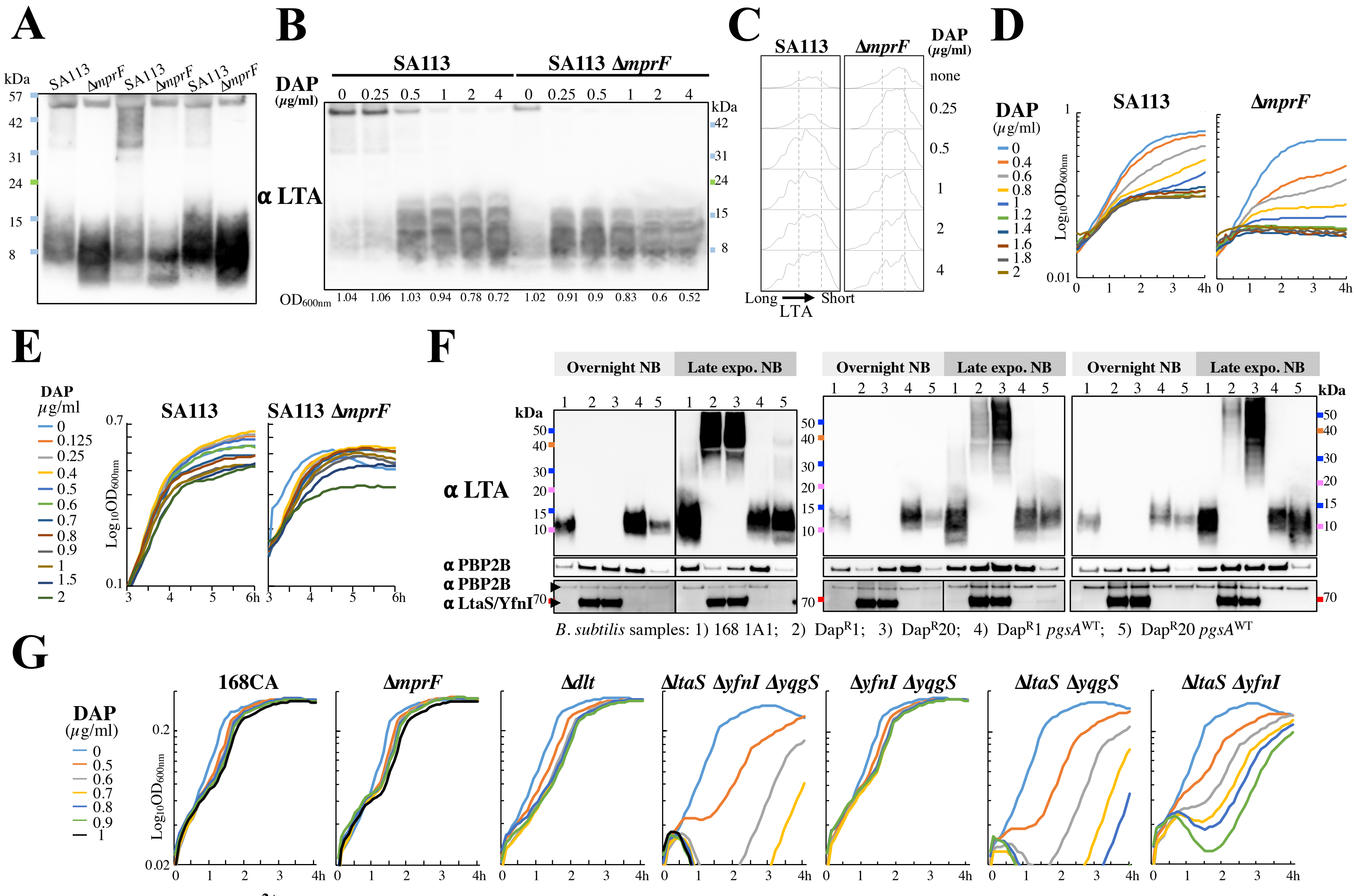


**Figure S10. Effects of Ca$^{2+}$-daptomycin on growth and LTA production in *B. subtilis* and *S. aureus*.**

**Figure S10. Effects of $Ca^{2+}$-daptomycin on growth and LTA production in *B. subtilis* and *S. aureus.***

**(A)** Samples of methicillin-sensitive *Staphylococcus aureus* strains SA113 and SA113 *ΔmprF* (1) were probed for LTA by Western Blot. Three independents set of cultures grown in LB at 37°C were tested simultaneously.

**(B)** *S. aureus* SA113 and SA113 *ΔmprF* strains were initially grown exponentially for 3 hours, after which calcium-daptomycin (DAP) or calcium only (1.25mM final) was added to the culture for an hour. Samples collected were analysed by LTA Western Blot as presented in (**A**).

**(C)** Analysis of the blot image obtained in (B) processed in Fiji (ImageJ) using the *gel* analysis function to extract the LTA signal for each sample.

**(D)** The growth of strains SA113 and SA113 *ΔmprF* in presence of calcium-daptomycin was monitored using a plate reader. Here cells were diluted at $OD_{600nm}$ of 0.05 in LB using a preculture of cells in exponential growth. Graphs represent an average of triplicate values.

**(E)** Growth of strains exposed to a range of calcium-daptomycin concentrations similar to that used in (**B**) when LTA changes were observed. Growth was monitored using a plate reader. Graphs represent the average of triplicate values.

**(F)** Detection of LTA in daptomycin resistant strains of *B. subtilis*. *B. subtilis* $Dap^R1$ and $Dap^R20$ mutants, carrying the *pgsA*(*A64V*) allele conferring daptomycin resistance (2), and strains 168 1A1 (wild type, WT), HB15516 ($Dap^R1$ $pgsA^{WT}$) and HB15507 ($Dap^R20$ $pgsA^{WT}$) were grown in NB overnight at 30°C and at late exponential growth phase (37°C, using theses overnight cultures). For each set of samples, LTA was detected along with PBP2B productions by Western blots. Bottom panel shows the cellular abundance of PB2B and the LTA-synthase detected with anti-LtaS antibody (that cross reacts with YfnI, unpublished Errington's lab). Samples used in the bottom panel where those of the LTA/PBP2B experiment loaded onto a new SDS-gel, after sample normalization based on PBP2B signal shown on the upper panel.

**(G)** Growth curves for *B. subtilis* 168CA, *ΔmprF* (AG1663), *Δdlt* (DLT71-CA), *ΔltaS ΔyfnI ΔyqgS* (AG600), *ΔyqgS ΔyfnI* (AG595), *ΔltaS ΔyqgS* (AG593), *ΔltaS ΔyfnI* (AG594) in NB at 37°C in presence of calcium-daptomycin and monitored using a plate reader. Graphs are representative of one set out of 3 independent experiments.
